# Unravel the Mystery of *NIC1*-locus on Nicotine Biosynthesis Regulation in Tobacco

**DOI:** 10.1101/2020.07.04.187922

**Authors:** Xueyi Sui, He Xie, Zhijun Tong, Hongbo Zhang, Zhongbang Song, Yulong Gao, Lu Zhao, Wenzheng Li, Meiyun Li, Yong Li, Yongping Li, Bingwu Wang

## Abstract

**Background:** Nicotine biosynthesis is mainly regulated by jasmonate (JA) signaling cascade in *Nicotiana tabacum*. As an allotetraploid species, the regulation of nicotine biosynthesis has been genetically verified via two unlinked *NIC* loci (named as *NIC1* and *NIC2*) which are possibly originated from its two ancestral diploids. Previously, a *N. tomentosiformis* originated ethylene response factor (*ERF*) gene cluster was identified as the *NIC2*-locus which has been demonstrated positively regulates nicotine accumulation in *N. tabacum*.

**Results:** Here, we describe the genetic mapping of *NIC1*-locus, the major nicotine regulatory locus, by using a *NIC1*-locus segregating population through bulked segregant analysis. We identified two linkage marker TM23004 and TM22038 were delimited the *NIC1*-locus within a ~34.3-Mb genomic region at pseudochromosome 07 of tobacco genome. Genomic scan within this region revealed a *NIC2*-*like* locus *ERF* gene cluster exist in. To verify this *ERF* gene cluster is the genetically called “*NIC1*-locus”, different functional experiments based on most of the *ERFs* in regulating nicotine biosynthesis and their influences on alkaloid accumulations have been carried out. Collinearity analysis showed that *NIC1*-locus *ERF* genes are originated from *N. sylvestris* and exclusively expressed in root tissues. In addition, transcriptomic results indicate that *NIC1*-locus *ERF* genes are coexpressed with the *NIC2*-locus *ERF* genes and other nicotine biosynthetic genes and regulators after JA induction. Furthermore, the suppressed expression of four *ERFs* of the *NIC1*-locus genes corresponding with decreased *NtPMT* and *NtQPT* expression in *NtMYC2*-RNAi lines indicates the selected *NIC1*-locus *ERFs* function in downstream of *NtMYC2* in the JA signaling cascades. In the meanwhile, the alkaloid levels are also determined by the amplitude of the four *ERF* gene expressions in both wild type and LA mutant. Additionally, *in vitro* binding assays, transient activation assays, and ectopic expression in transgenic plants demonstrate that these *ERF* genes are able to bind the GCC-box elements residing in the step-limiting gene promoters (such as *NtPMT2*, *NtQPT2*) and functional redundant but quantitatively transactivate nicotine biosynthetic gene expression. For *nic1*-locus mutation, two different sizes of deletions (*nic1-S* and *nic1-B*) were identified which occurred at the surrounding regions of the *NIC1*-locus gene cluster, which might disrupt, to some extent, chromosomal microenvironment and change gene expression around the deletion regions (including *NIC1*-locus *ERFs*), resulting in the decreased expression levels of *NIC1*-locus *ERFs* (such as *NtERF199*) and reduced alkaloid accumulation in the *nic1*-locus mutant.

**Conclusions:** Our findings not only provide insight in to the mechanism of the *NIC1*-locus *ERFs* in the regulatory network of nicotine biosynthesis, but also unraveled the theoretical basis of the *nic1*-locus mutation in low nicotine mutant. These functional verified *NIC1*-locus *ERF* genes can be further used as potential target(s) for ethyl methanesulfonate-based mutagenesis to manipulate nicotine level in tobacco variety in tobacco breeding program.

## Background

Tobacco (*Nicotiana tabacum* L.) belongs to the Solanaceae family and is an important economic crop in the world. Nicotine acts as the predominant alkaloid in tobacco plant for defense against herbivore and insect attack (Saitoh et al., 1985; Steppuhn et al., 2004). In commercial tobacco variety, nicotine biosynthesis and accumulation directly determine smoke quality and addiction (Davis and Nielsen, 1999; Shoji and Hashimoto, 2011a). Nicotine biosynthesis is exclusively occurred in tobacco roots, and then subsequently translocated through the xylem to the aerial parts of the plant, and accumulated in the leaf where it is secreted by trichomes (Dawson, 1941; Thurston et al., 1966; Hashimoto and Yamada, 1994). The formation of nicotine is completed by condensing two nitrogen-containing rings, pyrrolidine ring and pyridine ring. Pyrrolidine ring formation starts from ornithine or arginine catalyzed by *arginine decarboxylase* (*ADC*) or *ornithine decarboxylase* (*ODC*) to form putrescine. Putrescine is then converted to *N*-methylputrescine through *putrescine N-methyltransferase* (*PMT*). The *N*-methylputrescine is oxidized and cyclized to form a methyl-Δ^1^-pyrrolium cation by *N-methylputrescine oxidase* (*MPO*) (Hibi et al., 1994; Burtin and Michael, 1997; Heim et al., 2007). Pyridine ring is derived from the derivative of nicotinic acid. Nicotinic acid is sourced from the nicotinamide adenine dinucleotide (NAD) biosynthetic pathway, and at least three enzymes participating in this metabolic conversion steps, including *aspartate oxidase* (*AO*), *quinolinc acid synthase* (*QS*), and *qunolinic acid phosphoribosyl transferase* (*QPT*) (Sinclair et al., 2000; Ryan et al., 2012). The condensation of the pyrrolidine and pyridine rings is mediated by two orphan oxidoreductase *A622* and berberine bridge enzyme-like proteins (*BBLs*) which lead to nicotine formation in tobacco roots (Hibi et al., 1994; De Boer et al., 2008; Kajikawa et al., 2009; Shoji and Hashimoto, 2011a). Newly synthesized nicotine is transported into vacuoles by two tonoplast-localized multidrug and toxic compound extrusion (*MATE*) family transporters, *MATE1* and *MATE2*, and then *nicotine uptake permease 1* (*NUP1*) is responsible for importing nicotine into cells (Shoji et al., 2009; Hildreth et al., 2011; Kato et al., 2015).

Topping or herbivorous damage could dramatically enhance nicotine biosynthesis in plant by triggering the JA signaling pathway (Baldwin, 1989; Shoji et al., 2008). JA singling cascade has been demonstrated extensively involving in diverse plant specialized metabolite biosynthetic processes, including nicotine biosynthesis, through regulating various transcriptional regulators (Kazan and Manners, 2008 & 2013; Shoji et al., 2010; Zhang et al., 2012; Zhou and Memelink, 2016). In *Arabidopsis*, *bHLH* (basic Helix-Loop-Helix) type transcription factor *AtMYC2*, acting as a master regulator in JA signaling cascade, was able to directly or indirectly integrate different phytohormone signal pathways, and function in diverse aspects of plant growth and development, such as lateral root and adventitious root formation, flowering time (Kazan and Manners, 2008 & 2013). Previously, transcriptomic analysis of both topping treated root tissue and JA treated BY2 cell lines demonstrated that extensive gene expression reprogramming of the entire tobacco secondary metabolic system was occurred, and hundreds of differential expressed genes are identified (Goossens et al., 2003; Yang et al., 2017; Qin et al., 2020). Half decades ago, genetic studies revealed that alkaloid content in tobacco is regulated by two unlinked genetic loci nomenclature *NIC1* and *NIC2*, respectively (Legg et al., 1969 and 1971; Shoji et al., 2010). The *NIC1* and *NIC2* loci act in a semi-dominant fashion and have dose-dependent effects on total alkaloid levels. Comparing with *NIC2*-locus, *NIC1*-locus shows 2.4 times stronger effects in nicotine accumulation (Legg et al., 1971). The identification of *NIC2*-locus is achieved through genomic comparison among different *NIC* genomic background mutants. The *NIC2*-locus is composed of at least 7 ethylene response factor genes (*ERFs*) which is able to quantitatively but not equivalently activate the expression of nicotine biosynthetic genes (Shoji et al., 2010). Further evidence demonstrates that the *NIC2* mutation is caused by chromosomal deletion in the allele where *NIC2*-locus *ERFs* located in (Kajikawa et al., 2017a). Among these *ERFs*, *NtERF189* acts as a major activator and with abilities to activate nicotine biosynthesis via directly targeting the GCC-box element of the committed enzymatic gene promoters (Shoji et al., 2010). On the other hand, NtMYC2 not only collaborates with NtERF189 to intensively activate nicotine pathway gene expression by targeting different *cis*-elements in their promoters, but also trans-activates the *NIC2*-locus *ERF* expression simultaneously (Shoji and Harshimoto, 2011b). Transgenic experiments showed that the *NtERF* genes belong to the subgroup IX were able to modulate nicotine accumulation in an overlapped but distinct manner. For instance, *NtERF32*, a non-*NIC2*-locus *ERF*, is able to manipulate nicotine pathway gene expression and control nicotine content levels in transgenic plant (Sears et al., 2014). In addition, ectopic expression of NtERF91 could activate nicotine pathway gene expression and enhance alkaloid accumulations (Sui et al., 2019). Another two NtERF proteins, NtERF221/NtORC1 and NtERF10/NtJAP1 could individually or collaborate with each other to activate *NtPMT* gene expression (De Sutter et al., 2005).

Recently, genome-wide studies revealed that gene duplication, transposable element insertion are two important evolutionary events which are essential to achieve root-specific nicotine biosynthesis in different *Nicotiana* species (Kajikawa et al., 2017a; Xu et al., 2017). Bioinformatic analysis revealed that a novel *NIC2-like*-locus *ERF* gene cluster which highly homologous to the *NIC2*-locus *ERFs* exists in tobacco genome (Kajikawa et al., 2017a; Rushton et al., 2008; Sui et al., 2019). However, whether this novel *ERF* gene cluster is the genetically called “*NIC1*-locus” and their influence on nicotine pathway gene expressions and the regulatory mechanism of alkaloid accumulation have not been thoroughly investigated.

In this work, bulked segregant analysis based marker screening identified that two *NIC1*-locus linkage markers was able to delimit the *NIC1*-locus within a ~34.3-Mb genomic region at pseudochromosome 07 of tobacco genome where was the *NIC2-like*-locus *ERF* gene cluster located in, suggesting this *ERF* gene cluster is a strong candidate of the *NIC1*-lcous allele. Bioinformatic and transcriptomic analysis suggested that these clustered *NIC1*-locus *ERF* genes are originated from *N. sylvestris*, root-specific expression, and coexpressed with *NIC2*-locus *ERF* genes, nicotine structural genes, and transcriptional regulators during JA induction. Most of these *NIC1*-locus *ERF* genes, if not all, are controlled by NtMYC2 in JA signaling cascade and their expression levels are highly correlated to nicotine levels in wild type variety and its low nicotine mutant. In addition, their binding targets and transactivation abilities on key nicotine biosynthetic pathway gene promoters (*NtPMT2*, *NtQPT2*) were also explored. Finally, we ascribe the *nic1*-locus mutation is mainly caused by chromosomal rearrangement based structural variation. Two different sizes of chromosomal deletions were identified at the very upstream of the *NIC1*-locus gene cluster which altered whole gene expression pattern around the deletion regions (including *NIC1*-locus *ERFs*), resulting in the decreased *NIC1*-locus *ERF* expression (such as *NtERF199*) in the *nic1*-locus mutant. We suggest these *NIC1-*locus *ERF* genes can be used as gene resources for ethyl methanesulfonate-based mutagenesis to manipulate nicotine level in tobacco breeding and variety development.

## Results

### Genetic mapping of *NIC1*-locus

Since the *NIC1*-locus was genetically independent with *NIC2*-locus in regulating nicotine accumulation in *N. tabacum* (Legg et al., 1969), we carried out genetic mapping of this locus using an *NIC1-*locus segregating population (F_2_) derived from the cross between high intermediate variety MAFC5 (HI, *AAbb*) and low alkaloid variety LAFC53 (LA, *aabb*), respectively. HI accumulates moderate nicotine (5.03 ± 1.10 mg/g), whereas LA accumulates with low nicotine (0.36 ± 0.114 mg/g) (**Fig. 1A**). Bulked segregant analysis were performed using DNA samples from two parental lines (HI and LA), F_1_ hybrid of HI × LA, lowest nicotine bulk (Bs_L, DNA samples from 15 plants with lowest nicotine content), and highest nicotine bulk (Bs_H, DNA samples from 15 plants with highest nicotine content) from the segregating population. Extensive marker screening was performed to identify linkage marker(s) co-separate with the *NIC1*-locus. In total, 13,645 simple sequence repeat (SSR) markers were screened and three markers (TM22038, TM23004, and TM22041) PCR products are linked with the *NIC1-*locus, albeit other non-specific amplifications occurred (**Fig. 1B**). These three linkage markers were used to genotype a total of 176 F_2_ population plants. Since the *NIC* locus showed a semi-dominant and dose-dependent effects on nicotine levels, it is difficult to precisely tell the genotype of each F_2_ population plants. To fix this issue, the genotypes of the F_2_ population plants were predicted by considering both nicotine contents and marker-based genotyping results. Using this strategy, the ratio of high nicotine (A_bb) to low nicotine (aabb) is 133:46 which statistically fits 3:1 (χ^2^ = 0.00187, df = 1, *P* = 0.90), indicating that inheritance of the *NIC1*-locus allele is according to a dominant fashion (**Fig. 1B**). Preliminary mapping using the F_2_ plants (*aabb*) enabled us to delimit the *NIC1-*locus within a ~34.3-Mb region at pseudochromosome 07 of tobacco genome (**Fig. 1C**; Xie et al., unpublished data). Therefore, a genomic survey was conducted to annotate all putative genes within the target region, taking advantage of the available tobacco genome sequence and annotation information. Within this ~34.3-Mb genomic region, a *NIC2*-*like ERF* gene cluster, encoding a member of the *ERF* type transcription factors, which are highly homologous to *NtERF189* and presumably function in regulating nicotine biosynthesis (Kajikawa et al., 2017a and 2017b; Rushton et al., 2008; Sui et al., 2019). Furthermore, single knockdown of *NtERF199* within this *ERF* gene cluster by CRISPR/Cas9-mediated genome editing in transgenic plants only accumulates ~50% alkaloid levels of the control in leaves (**Fig. 1D**). Collectively, this *ERF* gene cluster was considered as the *NIC1* allele (hereinafter named as *NIC1*-locus *ERF* genes). Functional characterizations of these clustered *ERF* genes were conducted as follows.

**Fig. 1.**
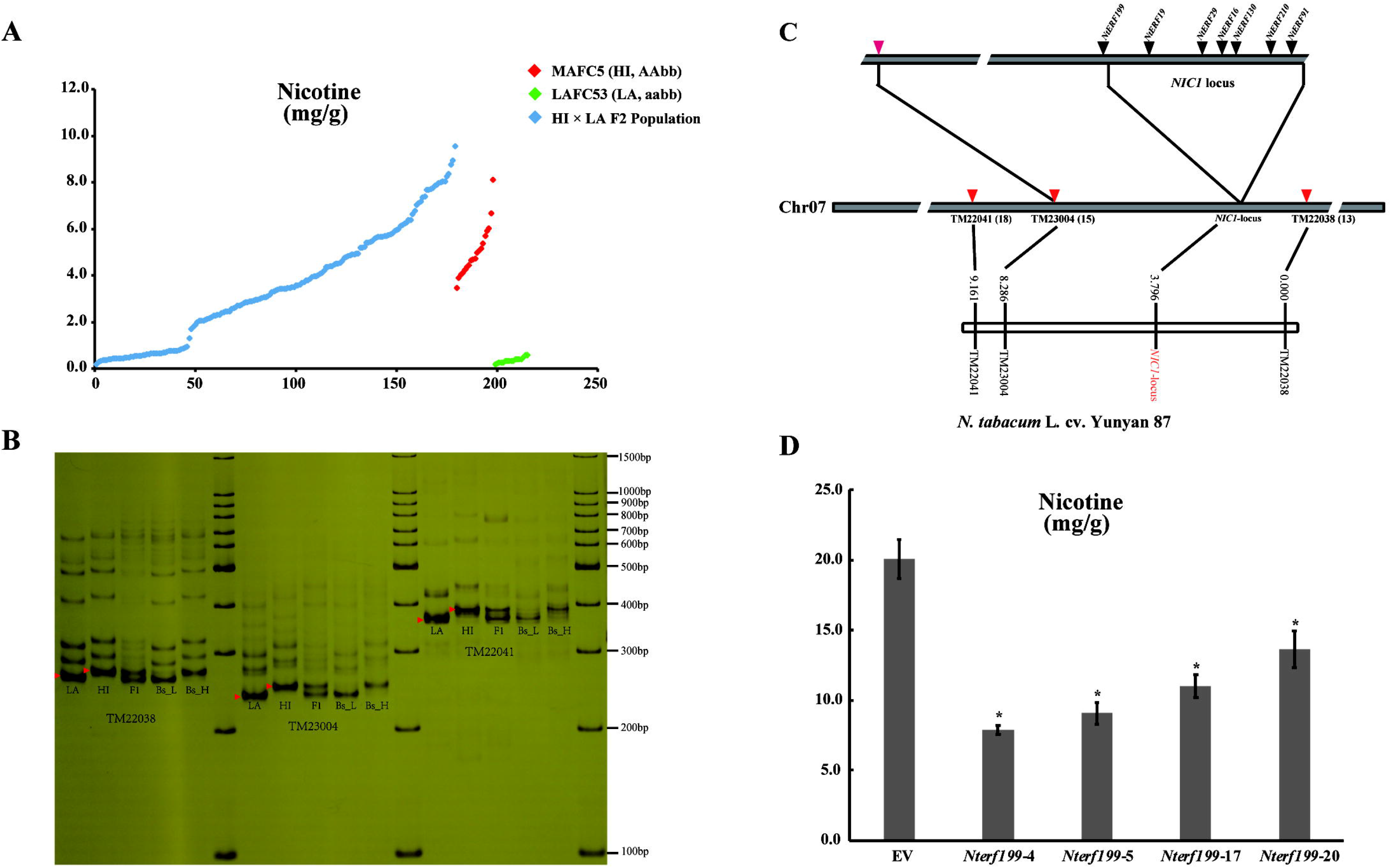
Genetic mapping of *NIC1-* locus. (A) Nicotine contents of two parental lines MAFC5 (*AAbb*, HI) and LAFC53 (*aabb*, LA) and their F_2_ population individuals (176 plants in total). (B) Three SSR markers TM22038, TM23004 and TM22041 were identified to link to the *NIC1-*locus. F_1_, hybrid of HI × LA; Bs_H: high nicotine bulk (15 highest nicotine contents individuals in the HI × LA F_2_ population) and Bs_L: low nicotine bulk (15 lowest nicotine contents individuals in the HI × LA F_2_ population). (C) Preliminary mapping result of *NIC1*-locus. The *NIC1-*locus was delimited to ~34.3-Mb genomic region between markers TM23004 and TM22038. Numbers in brackets indicate the number of recombination breakpoints separating the marker from *NIC1-*locus. The *NIC1*-locus genetic linkage map based on the F_2_ population was generated and the positions (cM) and names of markers in are denoted. Genomic scan of this ~34.3-Mb genomic DNA region within the pseudochromosome 07 of tobacco cv. Yunyan 87 (*AABB*) identified a *NIC2-like ERF* gene cluster, members of which are highly homologous to *NtERF189*, was considered as a strong candidate of the *NIC1* allele.

### Ancestral origin and root-specific expression of *NIC1*-locus *ERF* genes

Phylogenetic analysis of *NtERF* transcription factor family genes showed that both *NIC1*-locus *ERFs* and *NIC2*-locus *ERFs* were mutually clustered within the tobacco *ERF* family clad IX (**Fig. 2C**) (Rushton et al., 2008). Among these *ERFs*, *NIC2*-locus has proven was located in pseudochromosome 19 of tobacco genome and originated from *N. tomentosiformis* (Shoji et al., 2010). However, in comparison, the origin of the *NIC1*-locus *ERFs* remains unclear. To fix this issue, we firstly tried to clear the organization of tobacco genome based on T- and S-origins. Previously, synteny analysis clearly showed that intra-genome chromosomal rearrangement occurred between chromosomes of the *N. sylvestris* and *N. tomentosiformis* genomes, resulting in the synteny between Chr 07 and regions of both Chr19 and Chr14 in tobacco genome (Edwards et al., 2017). To further support this assumption, we performed genomic collinearity analysis by using nucleotide sequences of all *NIC1*-locus *ERFs* as queries to BLAST against *N. sylvestris* genome sequence in the China Tobacco Genome Database (https://10.6.0.76). As expect, the *NIC1*-locus *ERFs* appear to synteny with the *NsylERF* genes at the *Nsyl_Scf_33* scaffold of *N. sylvestris* genome (**Fig. 2A**, data not shown). Additionally, successful amplifications of all *NIC1*-locus *ERF* genes were obtained from both tobacco and *N. sylvestris* genomic DNA templates, and validated by sequencing (**Fig. 2B**) (Shoji et al., 2010). The expression pattern of all these *NIC1*-locus *ERFs* in different tobacco tissues was examined using qRT-PCR. These *NIC1*-locus *ERF* genes were preferentially expressed only in root tissues, whereas the expression of *NtERF16/NtERF210* was almost non-detectable in flower, mature leaves, stem in tobacco plant (**Fig. 3**).

**Fig. 2.**
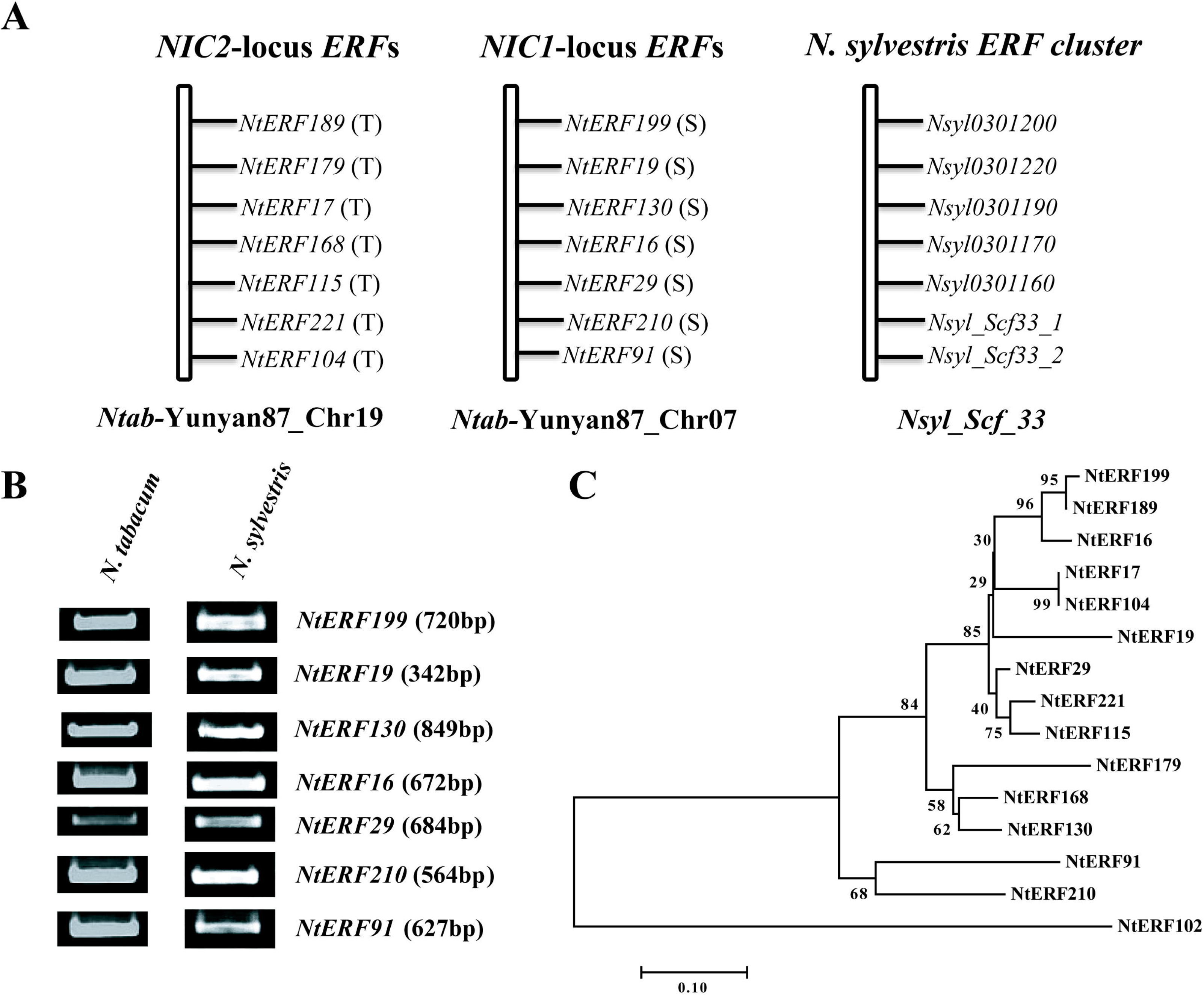
Collinearity analysis between *NIC2*-locus *ERF*s, *NIC1*-locus *ERF*s in *N. tabacum* genome and the ancestral *ERF* gene cluster in *N. sylvestris*. (**A**) Collinearity analysis of *NIC2*-locus *ERF* genes, *NIC1*-locus *ERF* genes in *N. tabacum* genome. Genomic scan within the genomic region between markers TM23004 and TM22038 identified a novel *ERF* gene cluster (*NIC1*-locus), which sharing highly amino acid similarities with *NIC2*-locus *ERF* genes (Ntab-Yunyan87-Chr19), located at Ntab-Yunyan87_Chr07 in the *N. tabacum* cv. Yunyan87 genome. The origin of all above mentioned *NtERFs* were indicated as T and S, respectively (T=*N. tomentosiformis*, S=*N. sylvestris*). To determine the genomic origin, the corresponding orthologues of the *NIC1*-locus *ERF* genes were identified in the Nsyl-Scf_33 scaffold of *N. sylvestris* genome by genomic collinearity analysis. (**B**) Genomic PCR analysis of *NIC1*-locus *ERF* genes. The genomic DNA from both *N. tabacum* cv. NC95 (HA, AABB) and *N. sylvestris* were successfully amplified using gene-specific primers and the fragment size of each *ERF* genes were presented in parentheses. (**C**) Phylogenetic analysis of seven *NIC2*-locus ERFs and *NIC1*-locus ERF amino acid sequences. A neighbor-joining phylogenetic tree was constructed using *NIC2*-locus *ERF* and *NIC1*-locus *ERF* protein sequences by the MEGA6 software. Bootstrap (2000 replications) analysis was performed to estimate the confidence of the topology of the consensus tree. Bootstrap support values are show above branches. NtERF102 protein sequence which belongs to the subfamily III of tobacco *ERF* family (Accession No.: XP_016504526) was used as an outgroup.

**Fig. 3.**
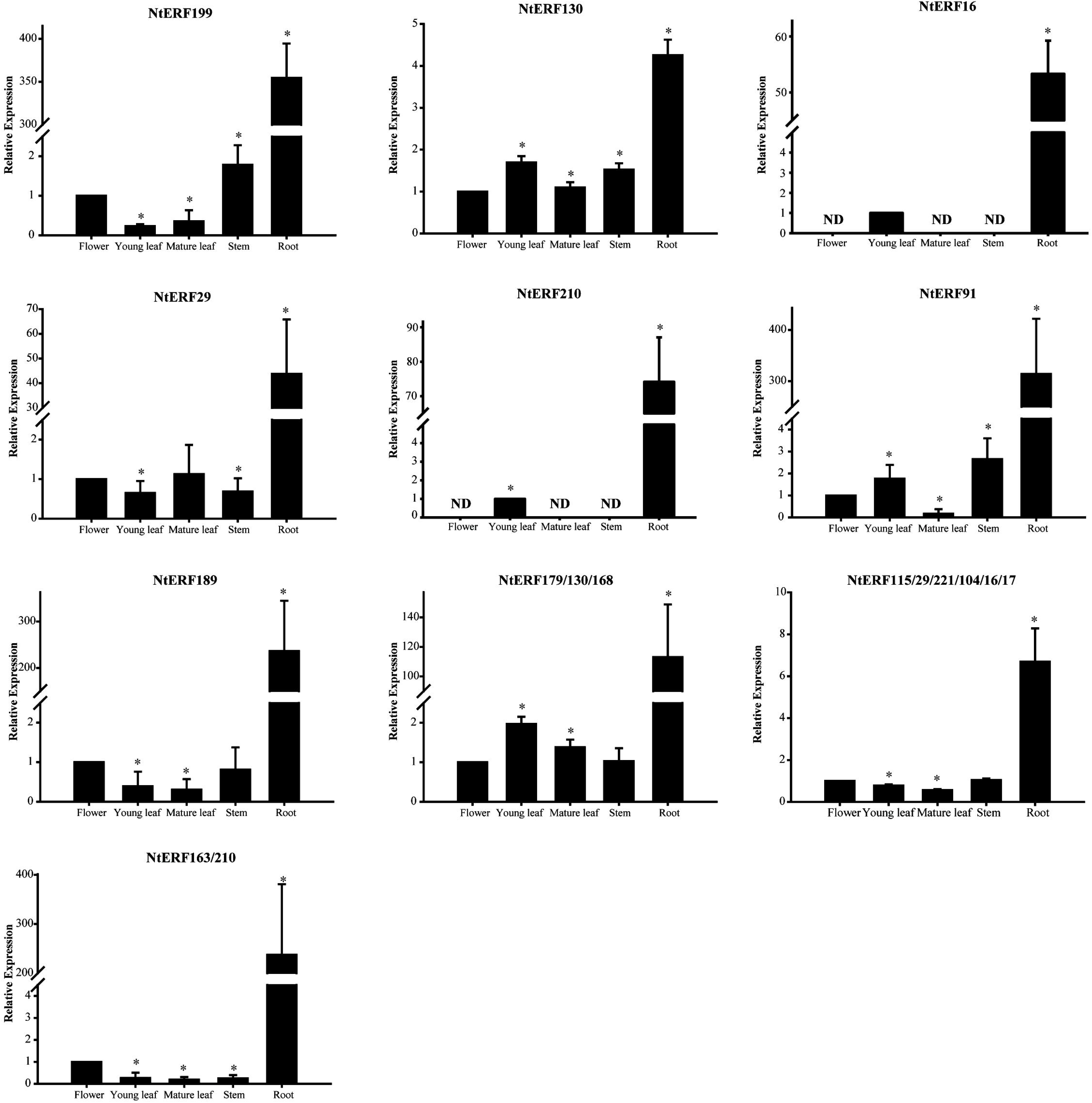
The tissue specific expression pattern of both *NIC1-* locus *ERF* and *NIC2*-locus *ERF* genes in different tissues of *N. tabacum*. Relative expression levels of the *NIC1-*locus *ERF* and NIC2-locus *ERF* genes in flower, young leaf, mature leaf, stem, and root in tobacco plants. Relative expression levels of *NIC1*-locus genes were determined by gene specific primer pairs, while most of *NIC2*-locus *ERF* expressions were determined by universal primer pairs using qRT-PCR. Tobacco actin gene (*NtActin*) was used as an internal control. Error bars represent SD (n=3). Asterisks indicate expressions that differ significantly (Student’s t-test, * P <0.05) from the flower tissue or young leaf (for *NtERF16* and *NtERF210* only).

### Expression of both *NIC1*-locus *ERFs* and *NIC2-*locus *ERFs* are highly correlated and response to JA induction

JA induced expression of structural genes and regulatory genes is a hallmark of the nicotine biosynthetic pathway (Zhang et al., 2012; Deway et al., 2013; Wang et al., 2015). The JA induced NtMYC2 and NtERF189 could transactivate nicotine pathway gene expression (such as *NtPMT2* and *NtQPT2*) in a synergistically manner (Shoji et al., 2010; Shoji and Hashimoto, 2012). To determine the correlation between the expression profile of all these *NIC1*-locus *ERF* genes with nicotine biosynthetic pathway gene expression, transcriptomes of MeJA treated tobacco root tissues were used. 0h MeJA treated roots were served as control. Hierarchical clustering analysis showed that *NIC1*-locus *ERF* genes (*NtERF199*, *NtERF19*, *NtERF210*, and *NtERF91*) were clustered with *NIC2*-locus *ERFs* (*NtERF115* and *NtERF221*), while *NtERF130/16/29* (*NIC1*-locus *ERFs*) were clade with *NtERF17/91L2/104/179*/168 (*NIC2-*locus *ERFs*) clad into another cluster, indicating all these *NtERF* TFs might involve in the same biological process, in this case, nicotine biosynthesis, to synergistically regulate downstream gene expression (**Fig. 4A**). On the other hand, expression of *NtMYC2a/b*, the regulatory hub of JA signaling cascades, were not only correlated with both *NIC1*-locus (such as *NtERF199*) and *NIC2*-locus (such as *NtERF189*) *ERFs*, but also with most of nicotine pathway genes, such as *AO*, *A622*, *QPT*, *QS*, *MPO*, *PMT2*, *ADC*, *MATE1/2*, *BBLa/b*, and *ODC*. These observations are consistent with the fact that *NIC1*-locus *ERF* TFs are JA inducible (**Fig. 4A**).

**Fig. 4.**
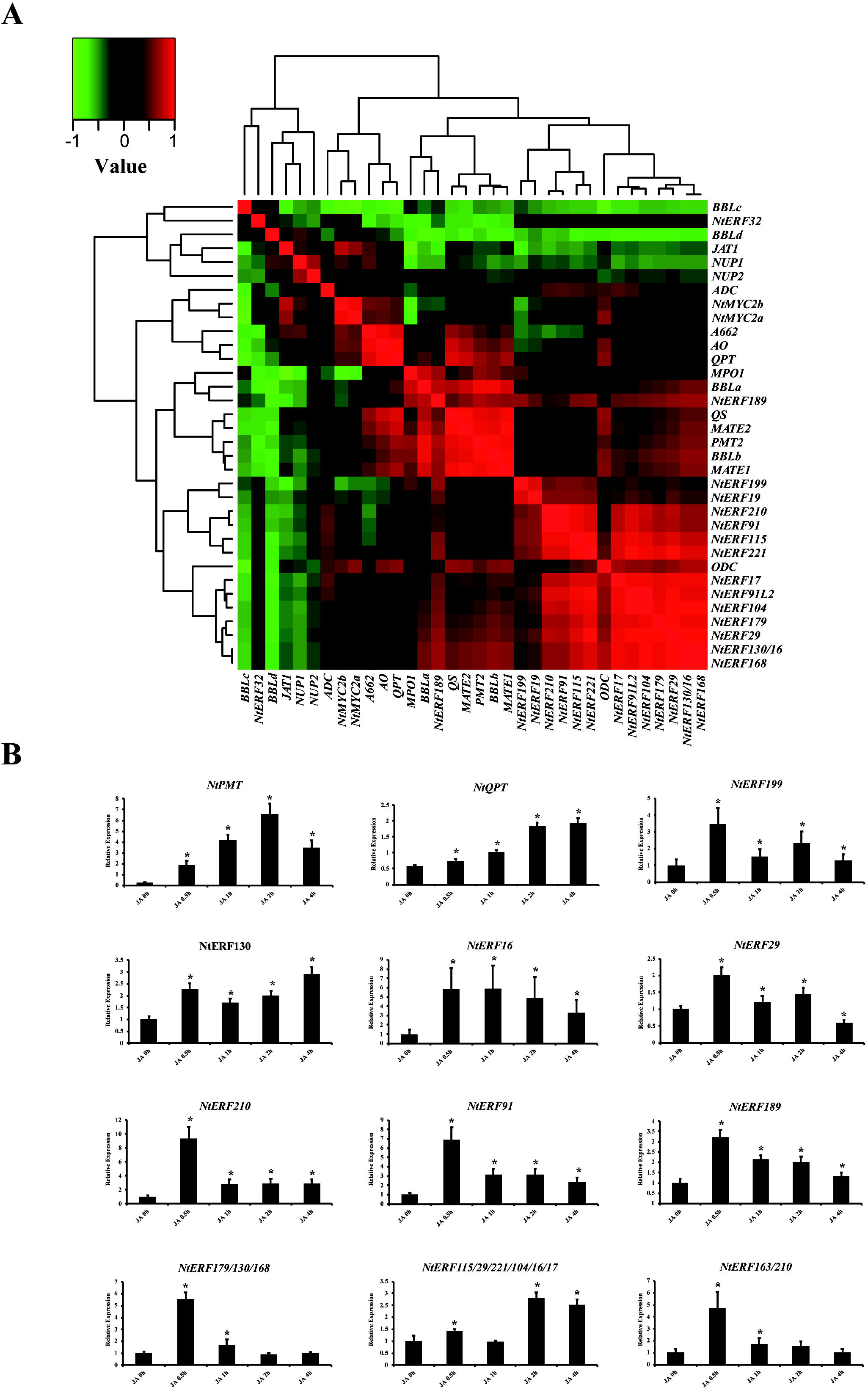
Correlation expression analysis of *NIC1*-locus *ERF* genes, *NIC2-* locus *ERF* genes together with nicotine biosynthetic pathway enzymatic and regulator genes using JA-treated tobacco root transcriptome. (**A**) Hierarchical clustering and heatmap showed that the *NIC1*-locus *ERF* genes are clustered with the most of the *NIC2-*locus *ERF* genes. *NtERF189* was co-expressed with nicotine pathway enzymes, such as *MPO1*, *BBLa/b*, *QS*, *MATE1/2*, and *PMT2*. In the meanwhile, *NtERF199* was co-expressed with most of *NIC1*-locus and *NIC2*-locus *ERFs*. The master regulator(s) of JA signaling cascade, *NtMYC2a*/*b*, were co-expressed with nicotine pathway genes, such as *AO*, *A622*, *QPT*, and *ADC*. (**B**) qRT-PCR validation of JA-treated root transcriptomic data for both *NIC1-*locus *ERF* genes and *NIC2*-locus *ERF* genes. Relative expression levels of the *NIC1*-locus *ERF* genes, nicotine biosynthetic pathway enzymes *NtPMT* and *NtQPT* were determined by gene-specific primer pairs, while most of *NIC2*-locus *ERF*s were determined by universal primer pairs. Tobacco seedling roots were treated with 100μM MeJA for different time points. Tobacco actin gene (*NtActin*) was used as an internal control. Error bars represent SD (n=3). Asterisks indicate expressions that differ significantly (Student’s t-test, * P<0.05) from 0h JA-treated root tissues.

To validate these transcriptome results, the transcript levels of all abovementioned *ERF* genes and two hallmark genes of nicotine pathway (*NtQPT* and *NtPMT*) were determined by qRT-PCR (**Fig. 4B**). As illustrated, the expression of all these *NIC1*-locus *ERF* genes was dramatically increased at various levels, particularly in 0.5h MeJA treated roots compared with the 0h control (**Fig. 4B**). Simultaneously, expression of the *NIC2*-locus *ERFs* also showed a similar expression pattern. In tobacco, *NtQPT2* and *NtPMT2* were known targets of the *NIC2*-locus NtERF189 TF in transcriptional regulation of nicotine biosynthesis (Shoji et al., 2010; Shoji and Hashimoto, 2011b and 2012). Consistent with these results, expression of *NtQPT* and *NtPMT* was gradually increased 2-6.8 and 0.5-2 folds and reached their peak expression at 2h, and 4h, respectively. This observation indicates the *NIC1*-locus *ERF* genes might function in the upstream of nicotine structural genes (such as *NtPMT* and *NtQPT*) and presumably by targeting the same *cis*-elements in their promoters as NtERF189 does.

### The *NIC1*-locus ERFs might function in downstream of NtMYC2 in JA signaling cascade

Acting as a multifaceted regulator, MYC2 has been proven involved in diverse plant growth and development processes. Specifically, MYC2 is the regulatory hub in JA signaling cascade and many MYC2 orthologs identified in different medicinal plant species were found positively regulating various plant specialized metabolite biosynthetic processes (Kazan and Manners, 2008 & 2013). In tobacco, RNAi-mediated suppression of *NtMYC2* significantly decreased expression levels of nicotine pathway enzymes such as *NtPMT1a*, *A662*, and *NIC2*-locus *ERF* TFs (Zhang et al., 2012; Shoji and Hashimoto, 2011b and 2012).

To confirm that *NIC1*-locus *ERF*s act downstream of NtMYC2 in JA signaling cascade, their expression levels were measured in *NtMYC2*-RNAi transgenic lines (**Additional file 1; Fig. S2**). Expression of *NtERF199* was decreased by 70% on average in *NtMYC2-*RNAi lines. Expression of four *NIC1*-locus genes, including *NtERF29*, *NtERF210*, and *NtERF91*, were decreased by 15%, 66%, and 48% on average, respectively, in *NtMYC2*-RNAi transgenic lines compared with empty vector (EV) control (**Fig. 5**). However, *NtMYC2* suppression failed to repress the expression of *NtERF130* and *NtERF16* (**Fig. 5**). In the meanwhile, expression of *NIC2*-lous *ERFs* was also decreased significantly at different levels. Consequently, expression of *NtPMT* and *NtQPT* was decreased by 90% and 73.5% on average, respectively, in *NtMYC2-*RNAi lines, compared to the EV control (**Fig. 5**). Taken together, these findings suggest that *NtMYC2* functional upstream of the *NIC1*-locus *ERFs* and nicotine pathway genes in JA signaling network.

**Fig. 5.**
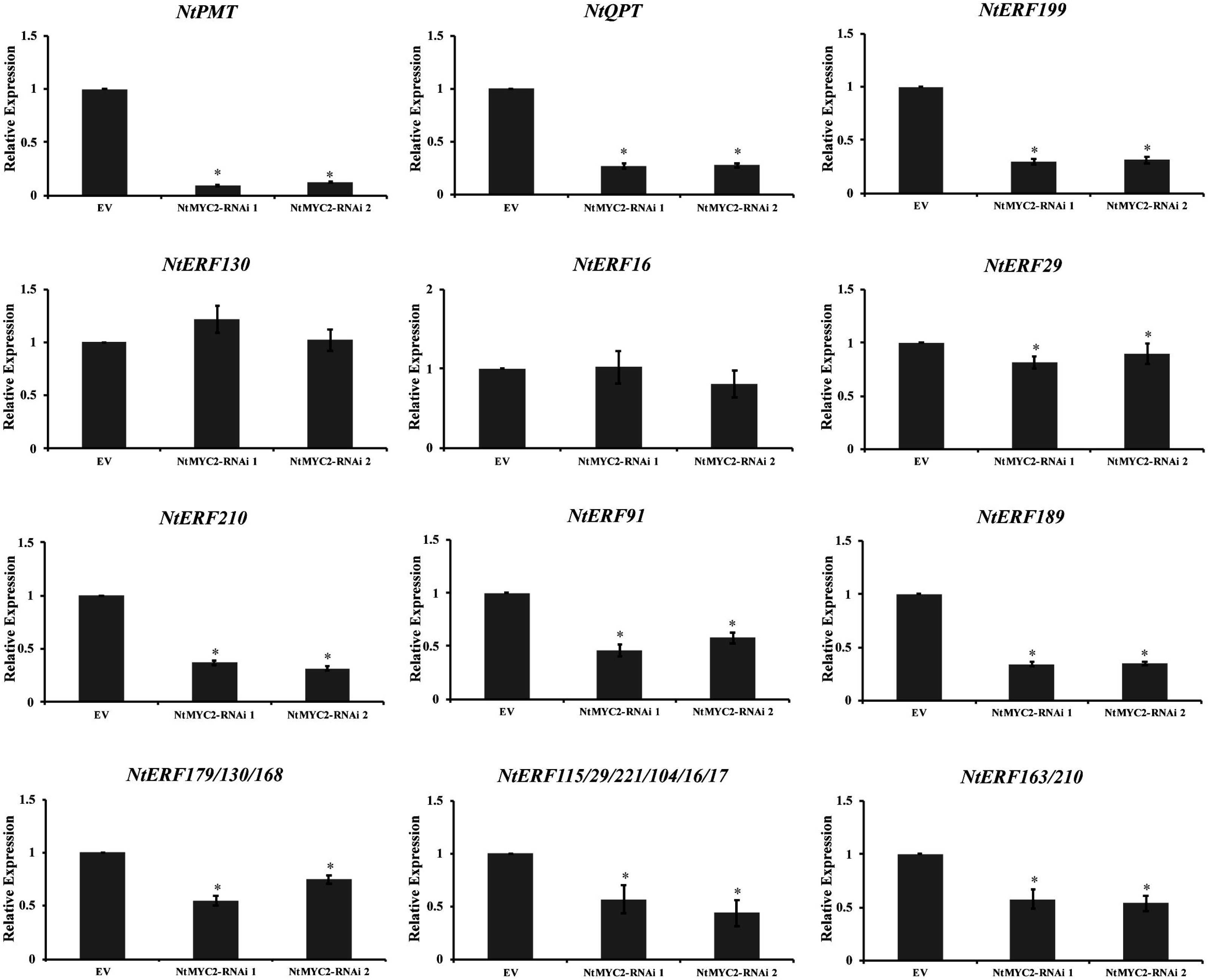
Relative expression of key nicotine pathway enzymatic genes, and both *NIC1*-locus *ERF* genes and *NIC2*-locus *ERF* genes in the root tissues of *NtMYC2-* RNAi transgenic lines. qRT-PCR validation of the expression levels of the *NIC1-*locus *ERF* genes in root tissues of the *NtMYC2*-RNAi transgenic lines. Relative expression levels of the *NIC1*-locus *ERF* genes, nicotine biosynthetic pathway enzymes *NtPMT* and *NtQPT* were determined by gene-specific primer pairs, while most of *NIC2*-locus *ERF*s were determined by universal primer pairs. Empty vector (EV) roots were used as control. Tobacco actin (*NtActin*) gene was used as an internal control. Error bars represent SD (n=3). Asterisks indicate expressions that differ significantly (Student’s t-test, * P <0.05) from EV root tissues.

### Expression levels of *NIC1*-locus *ERF*s are correlated with nicotine accumulation levels in tobacco

Accumulated evidences suggest the subgroup IX *NtERF* TFs could regulate multiple nicotine biosynthetic gene expression in an overlapped but divergent manner (Sears et al., 2014; De Boer et al., 2011; Sui et al., 2019). As a semi-dominant allele, the expression of *NIC1*-locus and *NIC2-* locus is strong associate with nicotine biosynthetic enzymes in tobacco (Saunders and Bush, 1979). To validate this assumption, we measured the expression levels of the *NIC1*-locus *ERF* genes and nicotine contents in HA lines and low nicotine mutant LA lines. As expected, the expression of these *NIC1*-locus *ERFs* was significantly compromised in LA mutants compared with HA wild type plants (except *NtERF130* and *NtERF16*). Expression of *NtPMT* and *NtQPT* was also decreased in LA mutants accordingly (**Fig. 6A**). In line with these results, HA plants produced 5.96 (mg/g), 202.23 (μg/g), 14.39 (μg/g) and 485.50 (μg/g) nicotine, nornicotine, anabasine, anatabine, respectively (**Fig. 6B**) (Reed et al., 2004; Cane et al., 2005). In contrast, LA plants showed significant decreasing in the accumulation of nicotine (0.87mg/g), nornicotine (88.32μg/g), anabasine (10.41μg/g) and anatabine (69.10μg/g) (**Fig. 6B**). Collectively, our findings highlight the highly positive correlation between *NIC1*-locus *ERF* gene expression and nicotine pathway enzymes expression, resulting in different alkaloid levels in *N. tabacum*.

**Fig 6.**
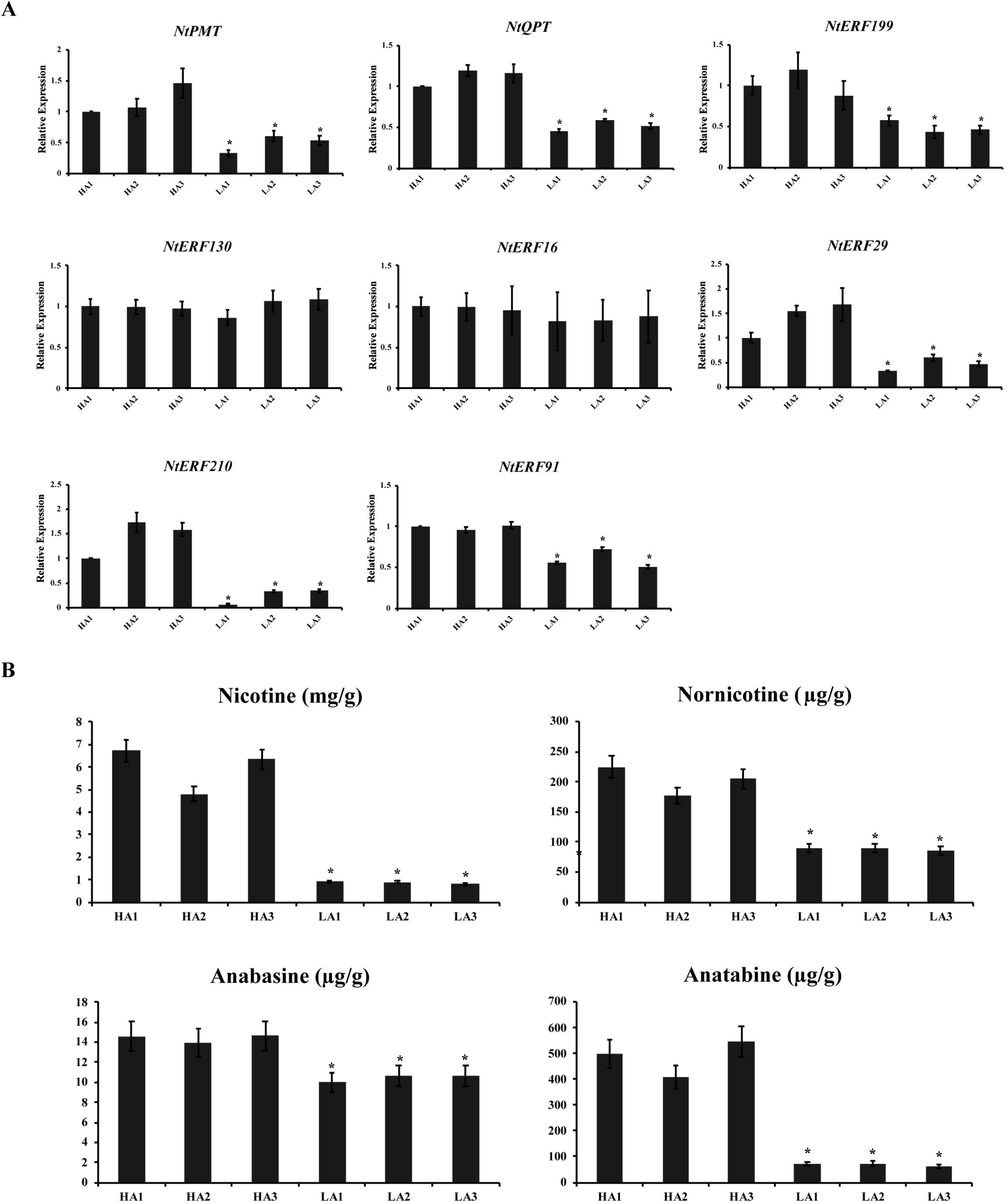
Relative expression analysis of *NIC1*-locus *ERF* genes and nicotine pathway enzyme genes and alkaloid profiles in both wild type HA and LA mutant lines. (**A**) qRT-PCR validation of the relative expression levels of the *NIC1-*locus *ERF* genes and nicotine pathway gene *NtPMT* and *NtQPT* in the root tissues of wild type variety and its nicotine mutant LA variety, respectively. Tobacco actin (*NtActin*) gene was used as an internal control. (**B**) Quantification of nicotine, nornicotine, anabasine and anatabine in wild type HA (AABB) and nicotine mutant LA (aabb). Equal amount of leaf sample was used for each variety for total alkaloid extractions. Alkaloid extracts from two near isogenic varieties were quantified by the Agilent HP 6890 GC-FID system. Error bars represent SD (n=3). Asterisks indicate expressions that differ significantly (Student’s t-test, * P <0.05) from HA.

### The *NIC1*-locus ERFs regulate *NtPMT2* and *NtQPT2* expression through specifically targeting the GCC-box element in their promoters

*NIC2*-locus ERFs, such as NtERF189 and NtERF179 are known to bind the GCC-box element in *NtPMT2* promoter to quantitively regulate nicotine synthesis (Shoji et al., 2010; Shoji and Hashimoto, 2011c). Since high amino acid similarity between NtERF189 and NtERF199, it is reasonable to speculate that the regulatory mechanism of *NIC1*-locus *ERF* proteins on nicotine biosynthesis acts in a similar manner. To test this hypothesis, the binding assays on the GCC-box in the *NtPMT2* and *NtQPT2* promoters were conducted. DNA fragments containing the GCC-box elements in the promoter of *NtPMT2*, or *NtQPT2* were used as probes. The results showed that the four selected recombinant *NIC1*-locus *ERF* proteins (NtERF199/130/16/29) were able to bind the *NtPMT2*, or *NtQPT2* promoter fragment via targeting the GCC-box elements, resulting in mobility shifts. However, the binding was abolished by adding the mutated probes (**Fig. 7A**). Taken NtERF199 as an example, formation of the DNA-protein complexes was gradually abolished when increasing amounts of unlabeled probes to the mixture as a cold competitor (**Fig. 7B**).

**Fig. 7.**
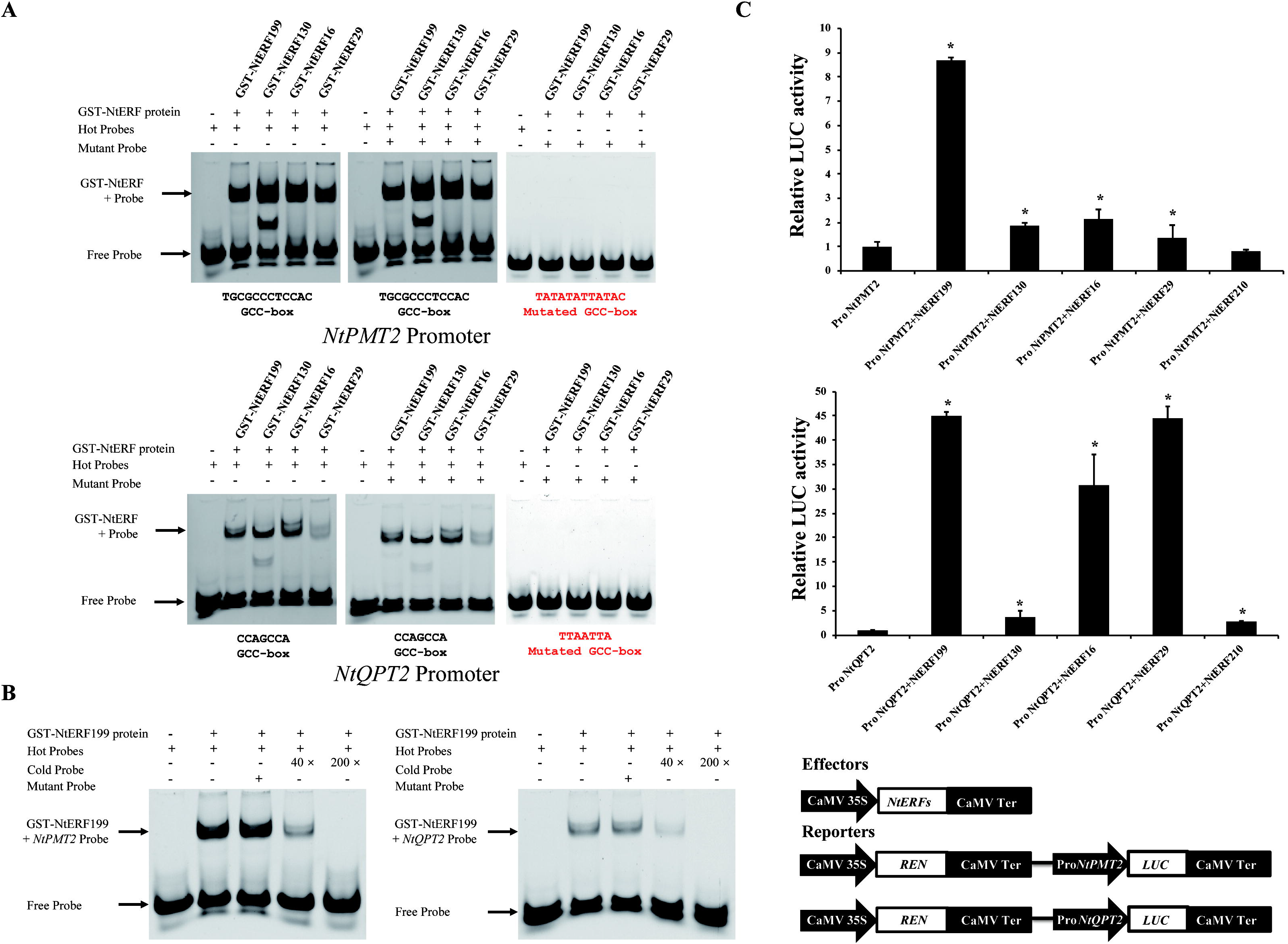
Analysis of binding abilities and transcriptional activities of the *NIC1*-locus *ERF* genes on the *NtPMT2* and *NtQPT2* promoters using electrophoretic mobility shift assay (EMSA) and transactivation assays. (**A**) *NIC1*-locus *ERF* genes were able to bind to the GCC-box within the *NtPMT2* and *NtQPT2* promoter. NtERF199, NtERF130, NtERF16, and NtERF29 were able to bind to the promoters of *NtPMT2*, and *NtQPT2* via directly targeting GCC-box element. Biotin-labeled DNA probe from the promoters or/and mutant probe was/were incubated with GST-NtERFs protein, and the DNA-protein complexes were separated on 8% native polyacrylamide gels. (**B**) The unlabeled (cold) probe consensus oligonucleotide was added at 40× and 200× fold excess over the labeled probe for competition and testing of binding specificity. (**C**) Transactivition transient assays demonstrate that the *NIC1-*locus ERFs differentially activated transcriptional activities of *NtPMT2* and *NtQPT2* promoters. *NtPMT2* and *NtQPT2* promoters fused to *luciferase* reporter were infiltrated into *N. benthamiana* leaves either alone or co-transformed with individual *NtERFs*. The reporter plasmid containing the *REN* reporter, controlled by the *CaMV* 35S promoter and *CaMV* 35S terminator, was used as a control for normalization. Luciferase activity was normalized against *REN* activity. The control represents the reporter alone without effectors. Error bars represent standard deviations. *P < 0.05(n = 3), by Student’s t-test.

To determine transactivation of *NIC1*-locus ERFs, *Agrobacterium*-infiltrated based transient luciferase assay was performed as well. Different combinations of *NIC1*-locus *ERFs* and the promoter regions of *NtPMT2* and *NtQPT2* (~1.5kb) as effector and reporter, respectively. The *NtPMT2*, or *NtQPT2* promoter, fused to the *firefly luciferase* (*LUC*) reporter gene, were infiltrated into *N. benthamiana* leaves alone or with plasmids expressing individual *NIC1*-locus *ERF* genes. As expected, NtERF199, NtERF130, NtERF16, or NtERF29 was able to activate the transcriptional activities of the *NtPMT2* promoter by 8.7, 1.8, 2.1, and 1.4- fold, respectively, compared to the control (**Fig. 7C**). Remarkably, NtERF199 exhibited strongest transactivation ability on the *NtPMT2* promoter. Similarly, different ERF proteins also showed various degrees of activation on *NtQPT2* promoter. As illustrated, the expression of *NtQPT2* promoter was dramatically increased when transactivated by NtERF199, NtERF16, NtERF6 proteins (30-45-fold higher). In addition, moderately activations were also observed on NtERF130 and NtERF210 proteins by 3.6-fold, and 2.7-fold, respectively (**Fig. 7C**). Taken together, these results indicate that these *NIC1*-locus ERF proteins might differentially activate key nicotine pathway enzymatic gene expression via targeting to the corresponding promoter regions *in vivo*.

### Ectopic expression of members of *NIC1*-locus *ERFs* quantitatively but not equivalently activate nicotine biosynthetic gene expression and enhance alkaloid accumulations in transgenic tobacco

Based on transient activation assay results, three *NIC1*-locus *ERFs* (*NtERF199*, *NtERF16*, and *NtERF29*) were selected to determine their individual contribution to alkaloid accumulations in tobacco. Transgenic tobacco lines overexpressing *NtERF199*, *NtERF16*, or *NtERF29* were generated. The relative transcript levels of nicotine biosynthetic pathway genes between the empty vector (EV) control and three independent transgenic lines of each *ERF* were measured by qRT-PCR (**Fig. 8A**). As illustrated, expression of *NtPMT* in *NtERF199*-OE lines was 1.8- to 3.3- fold higher than in the EV control line. Expression of the *NtODC* and *NtMPO* were increased by 2.5-, and 2.8- fold on average, respectively, in *NtERF199*-OE lines (**Fig. 8A**). Similarly, expression of the pyridine branch genes (*NtAO*, *NtQS*, and *NtQPT*) were also significantly increased by 2.3- fold, 1.5- fold, and 1.37- fold on average, respectively. This overall increased pattern of nicotine biosynthetic genes in *NtERF199*-OE lines (except for *NtADC*) indicates *NtERF199* acts as the major effector in *NIC1*-locus allele in regulating nicotine synthesis. Accordingly, similar coordinated expression pattern of nicotine biosynthetic genes was also obtained in both *NtERF16*-OE and *NtERF29*-OE transgenic lines. For *NtERF16*-OE transgenic lines, the relative expression levels of pyridine branch genes *NtAO* and *NtQPT* were significantly increased by 2.1-fold, and 1.3-fold on average, respectively. However, the relative transcript levels of *NtQS* were not significantly altered. In the pyrrolidine branch, expression of *NtODC* and *NtPMT* were upregulated by 3.1-fold, and 1.6-fold on average, and significantly increased expression of *NtADC* and *NtMPO* were only identified in *NtERF16*-OE 2 line (**Fig. 8A**). Additionally, expression of *NtAO* (2.3- fold), and *NtQPT* (1.5- fold) were significantly increased in the *NtERF29*-OE lines. On the other hand, the relative transcript levels of pyrrolidine genes *NtODC* (1.7- fold), *NtPMT* (1.9- fold), and *NtMPO* (2.0- fold) were also significantly upregulated at different levels, and significantly increased expression of *NtADC* was only identified in *NtERF29*-OE 4 line (**Fig. 8A**). Collectively, these results demonstrate that ectopic overexpression of these selected *NIC1*-locus *ERF* genes activate the expression of nicotine pathway genes in a functional redundant but distinct manner.

**Fig. 8.**
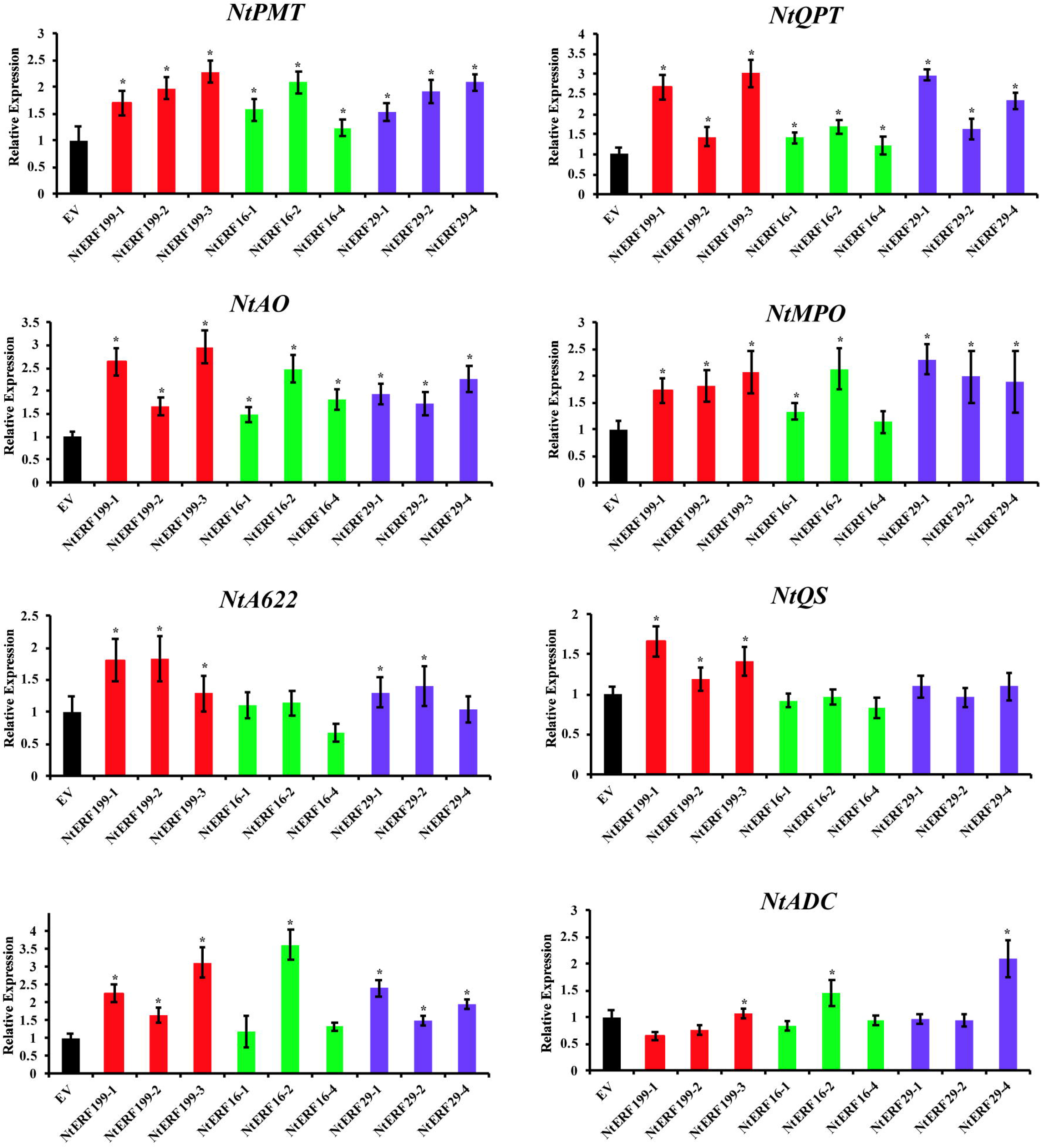
The expression pattern of nicotine pathway enzymatic enzymes and alkaloid accumulations in *NtERF199*, *NtERF16*, *NtERF29* overexpression lines. (**A**) Relative expression levels of the nicotine pathway enzymatic genes (*NtPMT*, *NtQPT*, *NtMPO*, *NtA622*, *NtQS*, *NtODC*, and *NtADC*) were determined by qRT-PCR. Tobacco actin gene (*NtActin*) was used as an internal control. Error bars represent SD (n=3). (**B**) Alkaloid accumulations in the *NtERF199*, *NtERF16*, *NtERF29* overexpression lines. Alkaloid extracts from the different leaves (tip, leave, lug) of the EV control and three OE lines for each *ERF* gene were analyzed, and the levels of nicotine, nornicotine, anabasine, and anatabine were measured. *P < 0.05(n=3), by Student’s t-test.

To overview metabolic consequences, alkaloid profile at different positional leaves (tip, leaf, and lug) of these *NIC1*-locus *ERF* transgenic plants were quantified. Among three *NIC1*-locus *ERFs*, *NtERF199*, highly homologous to *NIC2*-locus *NtERF189*, exhibits the strongest ability in promoting alkaloid accumulations in transgenic tobacco plants. Compared with the EV line, four alkaloids (nicotine, nornicotine, anabasine, and anatabine) were dramatically accumulated at various degrees from top to bottom leaves in *NtERF199*-OE lines. For instance, nicotine levels of the upper, middle, and bottom leaves in three transgenic lines were 2.18- fold, 1.89- fold, and 2.36- fold on average higher than the corresponding leaves of the EV control line (**Fig. 8B**). In comparison, the transactivation abilities of NtERF16 and NtERF29 in promoting nicotine biosynthetic gene expression were compromised under the same circumstance. Nicotine levels in the tip, leaf, and lug of three *NtERF16* (or *NtERF29*) transgenic lines were only marginally increased by 1.35 (or 1.31)- fold, 1.46 (or 1.34)- fold, and 1.22 (or 1.42)- fold on average, comparing with the EV control line. Furthermore, the stronger effect of NtERF199 than the other two *NIC1*-locus ERFs on enhancing nicotine biosynthesis could also be draw from the increasing trend of nornicotine and anatabine accumulations. In *NtERF199*-OE transgenic lines, nornicotine (or anatabine) levels were significantly increased by 5.53 (or 4.54)- fold, 2.33 (or 4.11)- fold, and 3.18 (or 4.77)- fold in upper, middle, and bottom leaves on average higher than in the EV control. Comparing with *NtERF199*, the increased fold changes of nornicotine of the upper, middle, and bottom leaves in *NtERF16*-OE (or *NtERF29*-OE) lines were 2.17 (or 2.21), 1.92 (or 2.25), and 2.09 (or 2.54) on average, respectively (**Fig. 8B**). In the meanwhile, the increased fold changes of anatabine of the upper, middle, and bottom leaves in *NtERF16*-OE (or *NtERF29*-OE) lines were 2.20 (or 2.58), 1.73 (or 2.75), and 1.62 (or 2.58) on average, respectively (**Fig. 8B**).

Even in the same type of leaves, contents of the four alkaloids in the *NtERF199*-OE lines were more pronounced than the other two *NIC1*-locus *ERFs* transgenic lines. Comparing with the EV lines, the four metabolites in the tips of the *NtERF199*-OE lines were significantly increased by 2.18- fold (nicotine), 3.53- fold (nornicotine), 1.63- fold (anabasine), and 4.52- fold (anatabine) on average, respectively. While these four metabolites in the tip leaves of the *NtERF16* (or *NtERF29*) transgenic lines were 1.35 (or 1.31)- fold, 2.17 (or 2.21)- fold, and 1.21 (or 0.98)-fold, and 2.20 (or 2.58)-fold on average higher than the EV control line. In line with this observation, similar accumulation trends of the four metabolites were also observed in middle, bottom leaves (**Fig. 8B**). Overall, the above data supported the notion that *NIC1*-locus ERFs act in a functional redundant but distinct manner and were able to quantitatively but not equivalently activate nicotine biosynthetic gene expression and metabolite accumulations, and *NtERF199* might be the dominant regulator within the *NIC1*-locus *ERFs*.

### Chromosomal deletions around the *NIC1*-locus region causing the decreased expression of *NIC1*-locus *ERFs* is the main reason for *nic1*-locus mutation in tobacco genome

In a previously study, the reasons for *nic2*-locus mutation in LA Burley 21 variety were well documented. A chromosome deletion occurred within the allele of *NIC2*-locus was identified (Shoji et al., 2010; Kajikawa et al., 2017a). To figure out the potential reasons for the decreased expression pattern of *NIC1*-locus *ERFs* in the *nic*1 mutant, genomic variation comparisons between HA and LA varieties were detected by whole genome resequencing analysis. The resequencing reads of two varieties were individually mapped back to the flued cured tobacco cv. Yunyan 87 genome. Consistent with previous results, previously reported *nic2* chromosomal deletion (~769 kb) was successfully identified. The identical *nic2* chromosomal deletion identified in the LA mutant indicates the resequencing data generated here were qualified to detect any potential structural variation, if exist, around the genomic region of the *NIC1*-locus on the pseudochromosome 07 of the LA genome. The genomic regions (~34.3-Mb, delimited by TM23004 and TM22038) where the *NIC1*-locus gene cluster located in were carefully examined. Surprisingly, two chromosomal deletions with different sizes were identified at the upstream of the *NIC1*-locus *ERF* gene cluster. As illustrated, a small deletion (~3 kb, hereinafter named as *nic1-S* deletion) was occurred at the very upstream of *NIC1*-locus *ERFs* (~ 40 kb distant from *NtERF199* promoter region); another large-scale (big) chromosomal deletion (~410 kb, hereinafter named as *nic1-B* deletion) was found located at the upstream of the *NIC1*-locus *ERFs* (~940 kb) based on the Yunyan 87 tobacco genome sequence. To verify these *in silico* identified deletions in *nic1* mutant, genomic PCR analysis was performed with specific primers. The specific primers were designed at different positions around both the *nic1-S* and *nic1-B* regions, donate as green arrowheads (**Fig. 9A**). Genomic DNAs from both HA and LA were used for the analysis. For the *nic1-S* deletion, successful amplifications designed for the 3 kb deletion (M2, 2022 bp) are obtained from HA but is missing in LA mutant. In contrast, another two specific amplifications (M1 and M3) designed at the genomic edges of the *nic1-S* deletion were successful amplificated (**Fig. 9A**). Same PCR strategy was also used for detecting the *nic1-B* deletion. There markers (M4, M5, and M13) designed to detect the genomic edges of the big deletion, and seven markers (M6–M12) which were evenly distributed within the *nic1-B* deletion region were used for confirming the deletion. As expected, successful amplifications were obtained from all primer pairs (M4–M13) by using the genomic DNA of HA as templates, while only three markers (M4, M5, and M13) with successful amplification fragments were detectable at the edges of the *nic1-B* deletion in LA mutant (**Fig. 9A**).

**Fig. 9.**
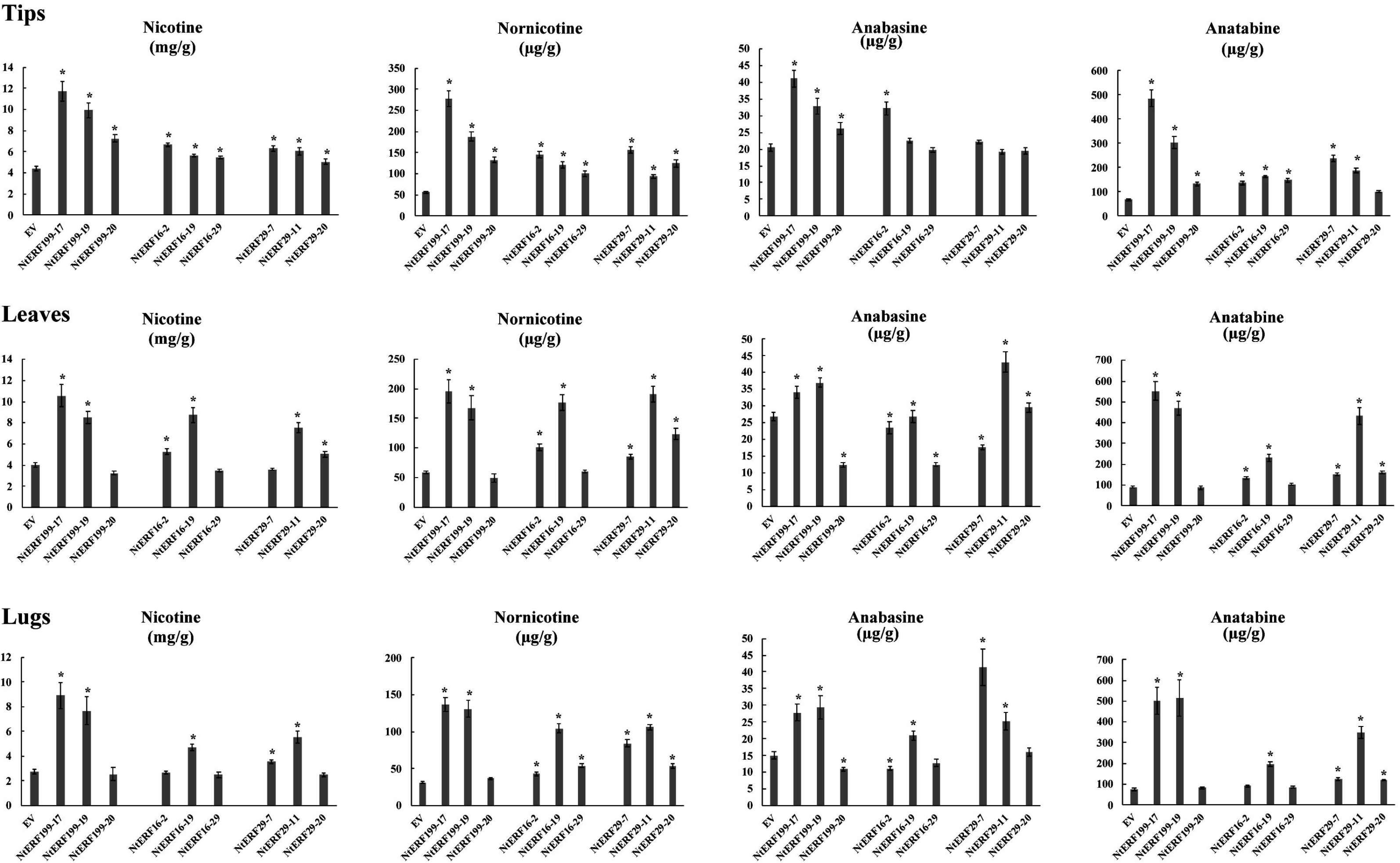
(**A**) The deletion model of *nic1*-locus in nicotine mutant line LA. Two different sizes of chromosomal deletions were found to located at the surrounding regions of the *NIC1*-locus *ERF* gene cluster (e.g. *NtERF199*) causing the *nic1*-locus mutant. Black arrowheads indicate the relative locations of *NIC1*-locus *ERFs*, while green arrowheads indicate the relative positions of specific markers (M1-M13) where *NIC1*-locus homozygous deletions which were present and absent in NC95 HA and LA varieties detected by using genomic PCR, respectively. (**B**) A regulatory model depicting *NIC1*-locus ERFs and *NIC2*-locus ERFs synergistically regulation of nicotine biosynthetic pathway in *N. tabacum*. *NtMYC2* acts as a regulatory hub that not only controls the *NIC2*-locus *ERF* TFs (e.g. *NtERF189*), but also simultaneously determines the expression levels of the *NIC1*-locus *ERF* TFs (e.g. *NtERF199*) TF genes to regulate nicotine pathway gene expression, such as *NtQPT* (pyridine branch), and *NtPMT* (pyrrolidine branch), respectively. To achieve quantitatively transactivation, two ancestral originated *NtERF* TF sets with differentially activation abilities are able to synergistically activating targeted gene expression by directly binding to the GCC-box motif within the promoter (such as *NtQPT*, *NtPMT*, and *NtQS*). In the meanwhile, *NtMYC2* also enhances the transactivity via physically interaction between NtMYC2 and the G-box containing promoters (such as *NtQPT*, *NtPMT*). Solid lines represent direct regulation and dashed lines represent regulation that may be direct or indirect, and lines with arrows represent transactivation.

To confirm the *nic1-S/B* deletions is responsible for the decreased expression of the *NIC1*-locus *ERFs* in the LA mutant, transcriptomic data of JA-treated root tissue from both HA and LA were used. In line with the results from previous studies, the FPKM value of *NtERF189* and *NtERF199* in the HA transcriptome are assigned with highest value, indicating that these two *ERFs* are major regulators in nicotine biosynthesis (Kajikawa et al., 2017a). In contrast, expression levels of *NIC1*-locus *ERFs* are dramatically decreased, even under detection limits, in LA mutant transcriptome (**Supplemental File 5**). We speculate that the *nic1-S* deletion (3kb), which located at only ~40 kb from the *NIC1*-locus *ERFs* (*NtERF199*) in tobacco genome, might be the essential reason for the decreased expression of these clustered genes, albeit more evidences are required for justify. Simultaneously, expression levels of other genes around both *nic1-S/B* deletion regions were also changed (upregulated or downregulated) in LA genetic background, comparing with its counterpart. Consequently, this fluctuant expression pattern caused by the *nic1-S/B* deletions might serve as a compensation to counterbalance the detrimental impact of chromosomal rearrangement on tobacco genome.

## Discussion

Genetic evidences have been demonstrated nicotine content in tobacco leaf is regulated by two independent loci *NIC1* and *NIC2* (Legg and Collins, 1971; Hibi et al., 1994; Shoji et al., 2010; Kajikawa et al., 2017a; Sui et al., 2019). Qualified genome sequences of different tobacco cultivars and their two ancestral diploids (Sierro et al., 2014; Edwards et al., 2017; Xie et al., unpublished data) provides us a chance to pinpoint the positions of segregating marker(s) and corroborate the genomic organization of the *NIC1*-locus *ERF* genes which is origin from *N. sylvestris* (**Fig. 1C; Fig. 2A**). The *NIC2*-locus comprises at least seven *AP2/ERFs* that target GCC-box elements residing in the promoter of *NsyPMT2 in vitro*. Suppression or dominant repression of the *NIC2*-locus *ERFs* moderately or severely reduced the expression of almost all structural genes in the pathway for nicotine synthesis (Shoji et al., 2010). Whereas overexpression of a single *NIC2*-locus *ERF* (*NtERF221*/*ORC1*) results in significantly increased nicotine accumulation (De Boer et al., 2011). As a typical regulator of the *NIC2*-locus *ERF* genes, the properties of NtERF189 in regulating nicotine biosynthesis have been extensively characterized. NtERF189 was shown to directly binds to the several nicotine structural gene promoters (such as *NtPMT2* and *NtQPT2*) and transactivate whole set of nicotine structural and transport gene expression (Shoji and Hashimoto, 2011b and 2011c). Overexpression or RNAi-mediated suppression of NtERF189 considerably increase or decrease the whole pathway gene expression levels and alkaloid contents (Shoji et al., 2010; Hayashi et al., 2020). MYC2 acts as master regulator in control of plant specialized metabolite biosynthesis, such as nicotine, terpenoid indole alkaloids, and artemisinin (Zhang et al., 2012; Zhang et al., 2011; Shen et al., 2016). In tobacco, NtMYC2 cooperates with NtERF189 to additively activate nicotine pathway gene expression via targeting the G-box elements within their promoters. Based on the observation that suppression of *NtMYC2* strongly co-repressed expression of most of nicotine structural and the *NIC2*-locus *ERF* genes, NtMYC2, presumably, is also able to control the expression of *NIC2*-locus *ERF* by directly binding to their promoters (Shoji and Hashimoto, 2011c and 2013).

*NtERF199*, acting as the homolog of *NtERF189*, have been predicted with similar function in nicotine biosynthesis regulatory network model (Shoji et al., 2010; Kajikawa et al., 2017b). Although both *NIC1*-locus and *NIC2*-locus *ERFs* are induced by jasmonates, *NtERF189* and *NtERF199* are only not response to salt stress (Shoji and Hashimoto, 2015). CRISPR/Cas9-mediated knock down of *NtERF189/NtERF199* dramatically reduces the expression of entire set of structural genes in the nicotine pathway and transport genes, together with low levels of alkaloid accumulation in transgenic tobacco lines (Hayashi et al., 2020). Nevertheless, whether this *N. sylvestris* originated *ERF* cluster, including *NtERF199*, is the so called “*NIC1*-locus” in genetic studies and the regulatory role of those *ERFs* in nicotine biosynthesis have not been thoroughly investigated.

We demonstrated that three SSR markers physically linked to the *NIC1*-locus allele and delimited the locus within a ~34.3 Mb region at the pseudochromosome 7 of tobacco genome (**Fig. 1C**). Genomic annotation of all putative genes within this region found a *NIC2*-*like* locus *ERF* gene cluster which was considered as strong candidate for the *NIC1*-locus (**Fig. 1A**). Synteny analysis showed that intra-chromosome rearrangement between tobacco ancestral diploids, resulting in high synteny between partial region of Chr19 and the entire region of Chr07, where were the positions of the *NIC2-*locus and *NIC2-like* locus (*NIC1*-locus) *ERF* cluster located in ((**Fig. 2A**, Kajikawa et al., 2017; Sui et al., 2019). In addition, the origin of the *NIC1*-locus *ERF* genes was confirmed by successful amplification of these genes in using DNA samples from *N. sylvestris* (**Fig. 2B**).

Accumulated evidences showed that the members of subgroup IX *NtERF* family in tobacco play functional overlapped but distinct roles in activating nicotine pathway genes (Ruston et al., 2008; Shoji et al., 2010; Todd et al., 2010; De Boer et al., 2011; Sears et al., 2014; Sui et al., 2019, Hayashi et al., 2020). Similar binding elements residing in these pathway structural gene promoters might serve as the prerequisite to achieve parallel transcriptional regulation (Kajikawa et al., 2017). In line with previously study, we found that the *NIC1*-locus ERFs specifically bound to GCC-box elements within the promoter regions of *NtPMT2* and *NtQPT2*, respectively (**Fig. 7B**). Transient overexpression of these *NIC1*-locus ERFs differentially activates the above-mentioned gene expression (**Fig. 7C**), suggesting that NtERF199 alone or in coordinate with the rest of ERFs synergistically modulates nicotine biosynthesis in tobacco. This functional divergence is supported by the amino acid differences in the *AP2/ERF* DNA-binding domain and /or the flanking regions of the *AP2/ERF* domain (Shoji and Hashimoto, 2012; Shoji et al., 2013). To consolidate this assumption, we found that the *NIC1-*locus *ERF199/16/29*, when individually overexpressed, has additive but not equivalent effects on the activation of nicotine pathway genes and the enhancement of alkaloid accumulations (**Fig. 8**). Similar to *NtERF189*, overexpression of *NtERF199* drastically enhancing alkaloid accumulation in tobacco through *in vivo* transactivation of the entire set of nicotine pathway genes. Whereas NtERF29 and NtERF16, due to limited trans-activity, only promotes expression part of those genes (**Fig. 8A**). Nevertheless, these case-by-case gene function justifications for both the *NIC1*-locus and *NIC2*-locus *ERFs* might not reflect the nature of the two *ERF* clusters in controlling nicotine biosynthesis *in vivo*. Since the *nic* mutations have does-dependent effects on nicotine levels, the nicotine contents of the *NIC1*-locus segregating population were highly plastic and versatile (**Fig. 1A**). Collectively, these observations imply that the ultimate alkaloid levels in tobacco might be depend on the transactivation activity outcomes powered by the intracluster interactions from single *NIC*-locus and / or intercluster interactions across the *NIC1*-locus *ERFs* and the *NIC2*-locus *ERFs*. Indeed, a recent study has shown that NtERF189 was able to activate the promoters of *NtERF115* and *NtERF179* within the *NIC2*-locus cluster (Paul et al., 2020). We thus propose that a similar scenario for intracluster manipulation within the *NIC1*-locus *ERFs*, to some extent, might be initially triggered by the transactivation of NtERF199, albeit additional experimental evidences are required.

This raises an intriguing question is why these *ERF* TFs are clustered and what is the benefit for nicotine production in tobacco. The plant metabolic clusters have been analyzed so far have shown this clustered architecture that confers complex trait maintenance, avoiding toxic intermediate accumulation, coregulation of gene expression, formation of metabolons (Boycheva et al., 2014; Nützmann et al., 2016). Root-specific expression of JA-responsive structural and regulatory genes is a hallmark of the nicotine pathway (**Fig. 3, 4**). Similar TF binding sites (such as GCC-box, G-box) residing in the structural gene promoters are important for their concerted transcription for nicotine biosynthesis (**Fig. 5, 6**). Further, genomic-wide studies revealed that the duplications of polyamine and NAD pathways and transposable element derived binding motifs in gene promoters forming a biosynthetic regulon which are essential for nicotine production efficiently in *Nicotiana* species (Shoji et al., 2010; Xu et al., 2017; Kajikawa et al., 2017). This coregulation based transcriptional activation of plant specialized metabolic pathway genes by jasmonate is likely conserved in plant. In *Catharanthus roseus*, a *bHLH* transcription factor iridoid synthesis 1 (BIS1) activates the promoters of all enzymatic genes that catalyze geranyl diphosphate to the iridoid loganic acid in terpenoid indole alkaloid (TIA) pathway (Moerkercke et al., 2015). The advantage of this parallel regulation is expression of an entire pathway with consequent avoid the accumulations of intermediate compounds may be toxic for plants (Boycheva et al., 2014; Nützmann et al., 2016).

Chromosome rearrangements (CRs) refers to chromosome abnormalities characterized by structural changes in chromosomes (Miller and Zachary, 2017). Based on whether net gain or loss of genetic material occurred, CRs have two forms: balanced (reciprocal translocations and inversions) and unbalanced (deletions and duplications) (Harewood and Fraser, 2014). These aforementioned chromosome structural variations have been shown to have far-reaching effects on gene expression. In general, the expression levels of genes situated within copy number variant (CNV) regions is positively correlated with gene copy number (Harewood and Fraser, 2014). The *nic2* genotype is caused by chromosomal deletion within the allele (Shoji et al., 2010; Kajikawa et al., 2017). Consistent with this, two *NIC* loci showed dose-dependent effects on nicotine levels and decreased expression pattern of nicotine pathway genes were observed in different *nic* genotype plants (**Fig. 6**; Shoji et al., 2010). Here, the nicotine contents of the *Nterf199* single mutants and *Nterf189/Nterf199* double mutants by CRISPR/Cas9-mediated genome editing prompted us to speculate that *NIC1*-locus *ERFs* and *NIC2*-locus *ERFs per se* are functional equivalent, regardless of chromosomal deletion effects on their expression (**Fig. 1D**; Hayashi et al., 2020). In this study, we showed that two different sizes of chromosomal deletions (*nic1-S* and *nic1-B*) are occurred at positions surrounding the *NIC1*-locus region with consequent not only reducing expression of *NIC1*-locus *ERFs* located nearby the breakpoint, but also affecting other genes in the region of the further afield (**Fig. 9A**). A reasonable explanation for this phenomenon is that the chromosomal rearrangements disrupts these *cis*-regulatory elements and/or the chromatin structure alteration changes the transcriptional control of *NIC1*-locus *ERFs*.

A recent study found that metabolic gene clusters usually reside in high dynamic regions in plant genomes. The cluster-associated interactive domains are capable to create local microenvironments to prevent the disturbs from neighboring chromosome areas and complete the coordinated gene expression (Nützmann et al., 2020). Root-specific synthesis of nicotine also suggests that exclusive spatial conformation constituted by cluster surrounding interactive domains is required to maintain transcriptional active status. Two deletions around the *NIC1*-locus region may disrupt this unique chromosomal environment, resulting in suppressed expression of *NIC1*-locus *ERFs*, even have a broadly effect on genome-wide gene expression changes. This assumption is supported by the facts that pronounced agronomic defects (such as yield penalty, poor leaf quality) were observed in *nic1* and *nic2* mutants, respectively (Legg et al., 1970).

To our knowledge, we propose a regulatory model of nicotine biosynthesis in tobacco. Perception of MeJA elicited expression of the *NtMYC2* genes. The NtMYC2 presumably trans-activates both *NIC2*-locus *ERFs* (e.g. *NtERF189*) and *NIC1-*locus *ERF* genes (e.g. *NtERF199*) through targeting the *cis*-regulatory elements residing in their promoters (**Fig. 9B**) (Shoji and Hashimoto, 2011b; Dewey and Xie, 2013). The initially translated ERFs might be able to activate their intra- and/or inter-cluster ERF genes to regulate nicotine production in a quantitative manner. Finally, the ERFs collaborate with MYC2 to activate gene expression by bound to the different cognate *cis*-elements (such as GCC-box and G-box) in the nicotine pathway gene promoters (such as *NtPMT2*, *NtQPT2*).

In sum, the *NIC1-*locus composed by seven *ERF* genes was identified via genetic mapping and genome synteny analysis. The *NIC1-*locus *ERF* genes are highly homologous with *NIC2*-locus *ERF* TFs. The *NIC1*-locus *ERFs* are response to JA and coordinate with *NIC2*-locus *ERF* TFs in regulating nicotine biosynthetic pathway. The expression levels of these *NIC1*-*ERFs* are positively related to pathway gene expression and nicotine contents in low nicotine mutants. *In vivo* binding assay, transient activation assay, *in vivo* gene function verifications revealed the underlying regulatory mechanism of the *NIC1-*locus *ERF* genes in pathway gene expression. The chromosomal deletions occurred at the *NIC1*-locus surrounding region might cause intrinsic chromatin conformation change and effect on gene expression of the *NIC1 ERF* gene cluster and other genes in the region of the further afield. We suggest these *ERF* genes can be used as the potential target(s) for ethyl methanesulfonate-based mutagenesis to develop low nicotine content tobacco variety in tobacco breeding program.

## Materials and methods

### Field trail and genetic mapping

The F_2_ mapping populations were derived from the cross between MAFC5 (*AAbb*, HI) and LAFC53 (*aabb*, LA). Two parental lines and their segregating population plants (176 individual) were grown in the experimental field during 2018 at Yanhe, Yuxi. Young leaves were sampled from HI, LA, F_1_ (HI × LA) lines, and individual F_2_ lines (176 plants in total) were stored in −80°C freezer for genetic mapping in future. All seedlings were transplant at the beginning of May, field managements were conducted as routine field managements. In late August, the leaves of parental lines and F_2_ population plants were harvested after two-weeks after topping and flue cured. A sample of leaves from each individual plant were taken for alkaloid quantification.

Genomic DNA isolation was conducted using the modified cetyltrimethyl ammonium bromide (CTAB) method (Maguire *et al.* 1994). To identify markers linked to the *NIC1*-locus region, bulked segregant analysis (BSA) was performed. Two pooled DNA samples (high nicotine/low nicotine) derived from the F_2_ population were made to make sure the two resultant bulked DNA samples were only genetically different at the *NIC1-*locus region and are seemingly heterozygous and monomorphic for all other regions. Each bulk contains selected 15 individuals with the highest (or lowest) nicotine contents. Two DNA bulks were made by mixing equal amounts of DNA from the selected plants. In total, 13,645 simple sequence repeat (SSR) markers were screened for identifying the segregating markers in the *NIC1-*locus region. PCR of the SSR markers, PCR product separation and visualization were performed as described previously (Tong et al., 2016). Genotyping of the F_2_ population were conducted using the linkage markers and the Kosambi mapping function was used to transform the recombination frequency to genetic distance (cM). The linkage analysis was performed using the program JoinMap^®^ version 4.0 (Van Ooijen et al., 2006).

### Experimental materials, RNA isolation, cDNA synthesis and gene cloning

*Nicotiana tobacum* var. ‘Yunyan 87’, NC95 (*AABB*, HA), and LAFC53 mutant (*aabb*, LA) seeds were germinated in cultural media in plastic pots at 25°C and 80% humidity in the tissue culture room with continuous light. For JA treatment, seven to eight-leaf stage seedlings were gently washed free of cultural media and the root tissues were immersed in 100 μM MeJA solution for 0, 0.5, 1, 2, and 4h, respectively. The treated roots were frozen in liquid nitrogen and store at −80 °C for further use. Total RNA isolated from JA-treated roots using RNeasy Plant Mini Kit (Qiagen, USA) was used for cDNA synthesis by following user’s instruction manual. The full-length cDNA of the *NIC1-*locus *ERF* genes were amplified using PrimeScript First-Strand cDNA Synthesis Super Mix Kit (Takara, Japan) from the first-strand cDNA with gene-specific primers with suitable restriction enzyme sites (**Additional file 3: Table S2**). PCR reactions were performed in a total volume of 50 μL containing 200ng of cDNA, 5×Phusion HF reaction buffer 10μL, 10mM dNTP 1μL, 1μL of each primer and 2U Phusion^®^ High-Fidelity DNA Polymerase (NEB, USA) for 35 cycles (98 °C for 30s, 98 °C for 7s, 57 °C for 30 s, 72 °C for 30s). PCR products were analyzed on a 1.5% ethidium bromide-stained agarose gel and purified for ligation. PCR products were cloned into the PCR^®^-Blunt II-TOPO vector (ABI, USA) and confirmed by sequencing.

### cDNA synthesis and qRT-PCR

Quantitative real-time PCR (qRT-PCR) was used to measure transcripts levels of the *NIC1*-locus *ERF* genes, *NIC2*-locus *ERF* genes, and nicotine pathway enzyme genes (*NtPMT* and *NtQPT*) in JA-treated root tissues. For tissue-specific expression, RNA samples were also isolated from flower, young leaf, mature leaf, stem, and roots, and the flower was used as a control. The qRT-PCR reaction was performed on an ABI7500 system (Applied Biosystems, USA), and Ultra SYBR Mixture (With ROX) kit was used (Kangwei, Beijing) by following SYBR Green I instructions. Gene relative expression level was calculated by using 2^−ΔΔCt^method.To measure expression of each *NIC1*-locus *ERF* genes, two primers were designed based on 5’-or 3’-untranslated region sequences. Universal primer pairs were used for detecting the *NIC2*-locus *ERF* gene expressions (Shoji et al., 2010). All primers used for qRT-PCR are listed in **Additional file 2: Table S1**. The *N. tabacum* actin (*NtActin*) gene was used as an internal control. All qRT-PCRs were performed in triplicates.

### Computational analysis

The genomic sequences of tobacco *ERF* genes were obtained from previous publications (Rushton et al., 2008) and their ORFs were translated by the online tool ExPASy Proteomics tools. Promoter analysis for *NtPMT2* (Accession No.: AF126809) and *NtQPT2* (Accession No.: AJ748263) were performed by AtPAN (http://atpan.itps.ncku.edu.tw/). To confirm sequence information, the CDS of the *NIC1*-locus *ERF* cluster genes were sequenced and verified in both *N. tabacum* and *N. sylvestris* genomes by BLAST against China Tobacco Genome Database (https://10.6.0.76) and NCBI database (https://www.ncbi.nlm.nih.gov/), respectively. Protein sequence alignment was conducted by using Clustal 2.0 software with the default parameters, and phylogenetic analysis was constructed by MEGA6 (NJ method with 2000 replicates).

### Sequencing library preparation and Illumina sequencing

RNA sequencing libraries of time course JA-treated root samples were made using 2 μg of total RNA following TruSeq^®^ Stranded mRNA Library Preparation protocol (Illumina, USA). Equal amount libraries were pooled and sequenced using Illumina Novoseq platform. Deep sequencing was performed in triplicates for each line with paired-end run (150bp × 2). The data quality was checked at the Novogene Bioinformatics Technology Co. Ltd (NBT, Beijing) and sequencing reads were provided as in FASTq format. Demultiplexing, adapter and barcode sequences trimming were performed on the DESeq software (Anders and Huber, 2010). For whole-genome resequencing, young leaves were collected from 3 individual HA and LA plants and mixed equally for genomic DNA extraction. Paired-end sequencing libraries (150bp × 2) were constructed for the HA and LA samples following manufacturer’s instructions (Illumina, USA). The libraries were sequenced in Illumina Genome Analyzer ×10 platform by NBT, Beijing. The tobacco reference genome Yunyan 87 (Xie et al., unpublished data) and its annotation were used.

### Co-expression analysis

For co-expression analysis, the transcriptomic data of different time course JA-treated root tissues were used. The differential gene expression analysis was performed by NBT, Beijing. Total reads were mapped to the tobacco genome (http://solgenomics.net/) using Top Hat (2.0.9) software (Edwards et al., 2017). Differential expressed genes between the control and treatment root tissues were defined with the edgeR software, based on log2 fold change >1 and adjusted P value◻<◻0.05. Pair-wise Pearson correlation coefficients for each transcript were calculated using the RPKM. Matrix distances for expression heat-map were computed with Pearson correlations of gene expression values (RPKM) by heatmap.2 function of gplots (version 3.0.1) Bioconductor package in R (version 3.2.2) (Wickham, 2009).

### Gel mobility shift analysis

The *NIC1*-locus *ERF* genes were individually cloned into the *EcoR*I/*BamH*I sites of pGEX-4T-1 vector (Amersham) to generate a GST fusion protein. Each recombinant GST-tagged NtERF protein was expressed in *E. coli* Rosetta (DE3) cells by induction with 0.1 mM IPTG for 3 h at 37°C, and the GST fusion proteins were purified by Glutathione HiCap Matrix columns (Qiagen) and eluted using 50 mM Tris-HCl, pH 7.2, 150 mM NaCl, and 30 mM glutathione. For EMSA experiments, four DNA probes, with or without biotin labeling, were synthesized based on the GCC-box element of the *NtPMT2* and *NtQPT2* promoters: Probe *NtPMT2* (5’-CTCTATTATATCGAG<u>TTGCGCCCTCCAC</u>TCCTCGGTGTC-3’) and Probe *NtQPT2* (5’-GTTTGTAAGCACT<u>CCAGCCAT</u>GTTAATGGAGTGC-3’) are identical to the *NtPMT2* promoter region (–149 to –138 relative to the TSS) and the *NtQPT2* promoter region (–152 to – 120 relative to the TSS) with the GCC-box (underlined), respectively. Probe *mNtPMT2* (5’-CTCTATTATATCGAG<u>TTATATATTATAC</u>TCCTCGGTGTC-3’) and Probe *mNtQPT2* (5’-GTTTGTAAGCAC<u>TTTAATTA</u>TGTTAATGGAGTGC-3’) are identical to the corresponding native promoters except that the GCC-box core sequences were destroyed by mutation (underlined). Complementary oligonucleotides, biotin labeled at the 5’end of each strand, were synthesized by Sangon Biotech Company (Shanghai) and annealed to produce double-stranded probes for EMSA. For mobility shift assays, 2 μg of purified GST-tagged ERF proteins were incubated with 2 ng of labeled probe, 200 ng of poly(dI-dC) to minimize nonspecific interactions, 25 mM Hepes-KOH (pH 7.4), 50 mM KCl, 0.1 mM EDTA, 5 % glycerol and 0.5 mM DTT in a 30 μL reaction volume. The mixtures were incubated at room temperature (20-25°C) for 20 min and separated on an 8 % (w/v) polyacrylamide gel in 0.5×TBE buffer. The band shifts were detected by autoradiographed on a FLA-9000 system (FujiFilm, Japan).

### Plasmid construction and dual luciferase transient assay

The pGreen0800-LUC reporter plasmids used in the protoplast assay contain both *firefly luciferase* (*LUC*) and *renilla* (*REN*) coding sequences driven by *NtQPT2* and *NtPMT2* promoters (approximately 1.5kb) and 35S promoter, respectively. The full length cDNAs of the *NIC1-locus ERF* cluster genes were cloned into the pGreenII 62-SK effector vector driven by 35S promoter. All the primers used were listed in **Additional file 3: Table S2**. *Agrobacterium tumefaciens* strain GV3101 containing different combination of effector constructs (35S-*NtERFs*) and/or reporter constructs was adjusted to OD_600_ = 0.2 in 0.5×MS medium with 150◻μM acetosyringone and 10◻mM MES (pH 5.6). The mixture was incubated at shaker for 3 hours in room temperature. The effector and reporter mixture were infiltrated into 4-week-old *N. benthamiana* leaves and kept in the dark overnight. The leaves of *N. benthamiana* were collected after 3 days for dual-*LUC* assay were conducted as described previously (Hellens et al., 2005).

### Tobacco transformation

The plasmids were mobilized into *Agrobacterium tumefaciens* strain C58C1 using the freeze-thaw method (Weigel and Glazebrook, 2006). For RNAi-mediated suppression, a 335 bp fragment was amplified from *NtMYC2* (Accession No.:GQ859160) and cloned in both sense and antisense orientation in pK7GW1WG2 vector. For overexpression, three *NIC1*-locus *NtERF199/16/29* were PCR amplified from tobacco root tissues and cloned in pK2GW7 vector containing *CaMV* 35S promoter and terminator. The empty pK2GW7 vector was used as an EV control. For CRISPR/Cas9-mediated mutagenesis, the *NtERF199* gene was sequenced from HA for gRNA design. The gRNA 5’-GATTCAAGGAGATGAGAGT-3’ (3’-ACTCTCATCTCCTTGAATC-5’) was designed to target the specific region of *NtERF199*. The gRNA sequences were cloned in the binary transformation vector pHSE401 (Xing et al., 2014) and confirmed by sequencing. Transformation of tobacco seedlings with *A. tumefaciens* and generation of the transgenic plant were conducted as previously described (Wang et al., 2015).

### Alkaloid Extraction and Quantification

For alkaloid extraction, lyophilized leaves collected from parental lines (HA and LA), 176 F_2_ segregating population plants, *NtERF199*-CRISPR/Cas9 and *NtERF199/16/29*-OE and empty vector (EV) transgenic plants were ground. Equal amounts of each samples were immersed in 2 mL of 2N NaOH solution and swirled for sample moisten. 10 mL of methyl tertiary butyl ether (MTBE) containing 0.1062 g/mL of quinoline was then added and extracted for 2.5 h on a shaker followed by resting overnight to separate. The extracts (~ 1 mL of the top MTBE layer) were then analyzed using the Agilent HP 6890 GC-FID system for alkaloid quantifications as previously described (Wang et al., 2005).

## Additional files

**Additional file 1: Figure S1.** The relative expression levels of *NtMYC2* in empty vector (EV) control and two *NtMYC2*-RNAi transgenic lines.

**Additional file 2: Table S1** Primer sequences used for qRT-PCR in this study.

**Additional file 3: Table S2** Primer sequences used for gene and promoter cloning.

**Additional file 4: Table S3** Primer sequences of *NIC1*-locus linkage markers and deletion detecting genomic PCR in this study.

**Additional file 5: Table S4** Expression levels of genes around *NIC1*-locus regions in HA and LA nicotine mutant.

## Abbreviations

A622: a PIP family oxidoredutase for nicotine precursors condensation
ODC: ornithine decarboxylase
ADC: arginine decarboxylase
PMT: putrescine *N*-methyltransferase
MPO: *N*-methylputrescine oxidase
AO: aspartate oxidase
QS: quinolinic acid synthase
QPT: qunolinic acid phosphoribosyl transferase
SPDS: spermidine synthase
SAMS: *S*-adenosylmethionine synthase
SAMDC: S-adenosylmethione decarboxylase
PCR: Polymerase chain reaction
RNA-seq: RNA Sequencing
RPKM: Reads Per Kilobase per Million mapped reads
SSR: simple sequence repeat
qRT-PCR: Quantitative real-time PCR
CR: chromosome rearrangement
CNV: copy number variant

## Acknowledgements

The authors express our gratitude to Dr. JQ. Wu (Kunming Institute of Botany, China) for comments on this manuscript. This research was supported by the Project of Yunnan Tobacco Company (Contract No. 2018530000241005), and the National Science Foundation of China (Grant No. 31860069) to X.S.

## Author’s contributions

X.S., B.W., and H.Z. designed the study; X.S., H.X., Z.T., Z.S., Y.L., Y.G., L.Z., W.L. and M.L. performed the experiments; X.S., H.X., Z.S., L.Z., W.L., Y.G., Y.L., Y.P.L., B.X. analyzed the sequencing data; X.S. and B.W. wrote the paper.

## Figure legends

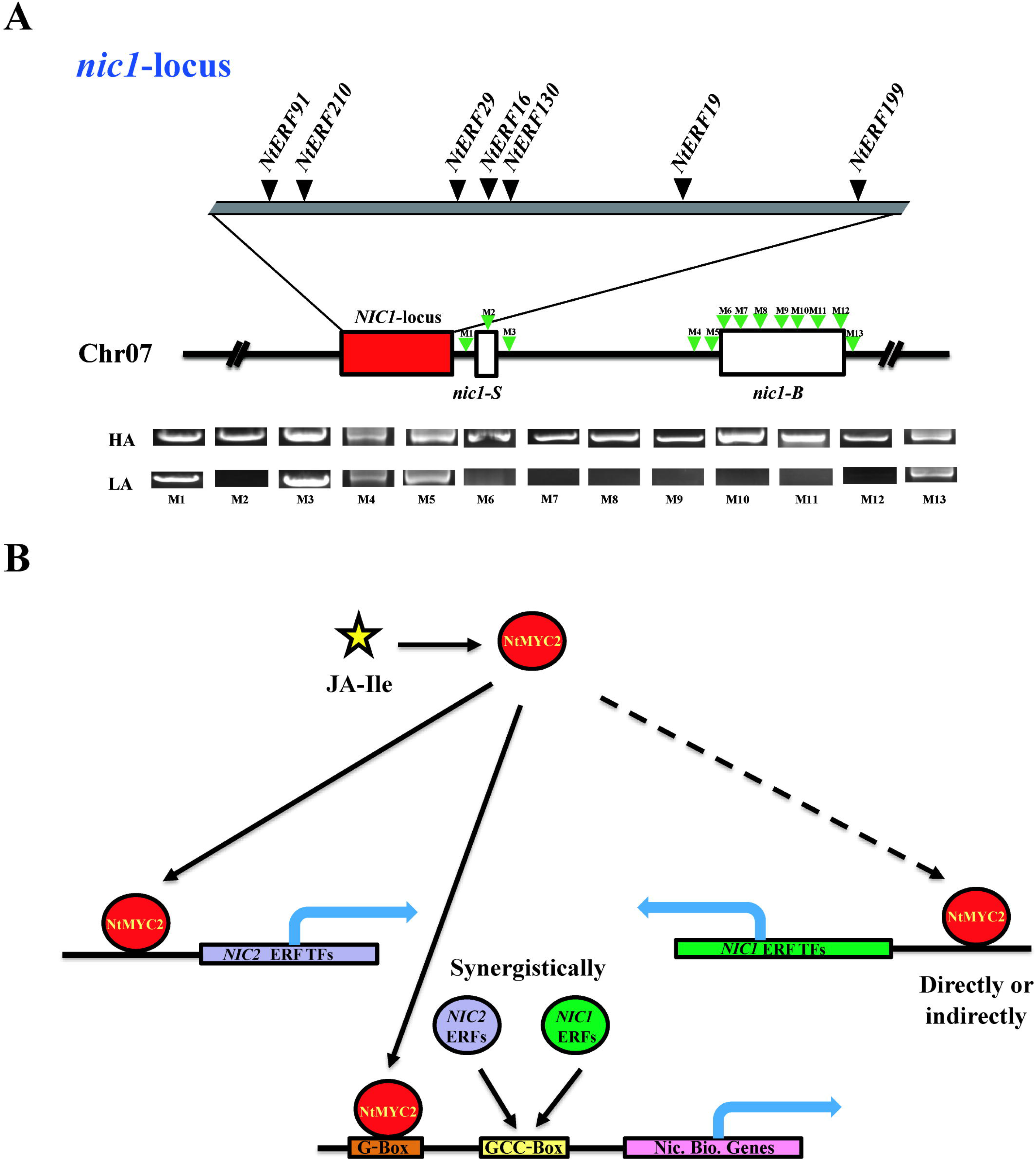

## References

Anders S, Huber W: Differential expression analysis for sequence count data. BMC Genome Biol. 2010, 11(10): R106.

Baldwin IT: Mechanism of damage-induced alkaloid production in wild tobacco. J. Chem. Ecol. 1989, 15(5):1661–1680.

Boycheva S, Daviet L, Wolfender J, Fitzpatrick TB: The rise of operon-like gene clusters in plants. Trends Plant Sci 2014, 19: 447–459.

Cane KA, Mayer M, Lidgett AJ, Michael A, Hamill JD: Molecular analysis of alkaloid metabolism in *AABB* v. *aabb* genotype *Nicotiana tabacum* in response to wounding of aerial tissues and methyl jasmonate treatment of cultured roots. Funct. Plant Biol. 2005, 32(4): 305–320.

Davis L, Nielsen M: Tobacco: Production, Chemistry and Technology. Oxford: Blackwell Science; 1999.

Dawson RF: The localization of the nicotine synthetic mechanism in the tobacco plant. Science 1941, 94(2443):396–397.

De Boer KD, Lye JC, Aitken CD, Su AK-K, Hamill JD: The A622 gene in *Nicotiana glauca* (tree tobacco): evidence for a functional role in pyridine alkaloid synthesis. Plant Mol. Biol. 2008, 69(3):299–312.

De Sutter V, Vanderhaeghen R, Tilleman S, Lammertyn F, Vanhoutte I, Karimi M, Inzé D, Goossens A, Hilson P: Exploration of jasmonate signalling via automated and standardized transient expression assays in tobacco cells. Plant J. 2005, 44(6):1065–1076.

Dewey RE, Xie J: Molecular genetics of alkaloid biosynthesis in *Nicotiana tabacum*. Phytochemistry 2013, 94:10–27.

Edwards KD, Fernandez-Pozo N, Drake-Stowe K, Humphry M, Evans AD, Bombarely A, Allen F, Hurst R, White B, Kernodle SP et al: A reference genome for *Nicotiana tabacum* enables map-based cloning of homeologous loci implicated in nitrogen utilization efficiency. BMC Genomics 2017, 18(1):448.

Goossens A, Hakkinen ST, Laakso I, Seppanen-Laakso T, Biondi S, De Sutter V, Lammertyn F, Nuutila AM, Soderlund H, Zabeau M et al: A functional genomics approach toward the understanding of secondary metabolism in plant cells. Proc Natl Acad Sci U S A. 2003, 100(14):8595–8600.

Harewood L, Fraser P: The impact of chromosomal rearrangements on regulation of gene expression. Hum. Mol. Genet. 2014, 23:76–82.

Hashimoto T, Yamada Y: Alkaloid biogenesis-molecular aspects. Annu Rev Plant Physiol Plant Mol Biol 1994, 45:257–285.

Hayashi S, Watanabe M, Kobayashi M, Tohge T, Hashimoto T, Shoji T: Genetic manipulation of transcriptional regulators alters nicotine biosynthesis in tobacco. Plant Cell Physiol. 2020; pcaa036.

Heim WG, Sykes KA, Hildreth SB, Sun J, Lu RH, Jeleko JG: Cloning and characterization of a *Nicotiana tabacum* methylputrescine oxidase transcript, Phytochemistry 2007, 68: 454e463.

Hellens RP, Allan AC, Friel EN, Bolitho K, Grafton K, Templeton MD, Karunairetnam S, Gleave AP, Laing WA: Transient expression vectors for functional genomics, quantification of promoter activity and RNA silencing in plants. Plant Methods 2005, 1:13.

Hibi N, Higashiguchi S, Hashimoto T, Yamada Y: Gene expression in tobacco low-nicotine mutants. Plant Cell 1994, 6(5):723–735.

Hildreth SB, Gehman EA, Yang H, Lu RH, Ritesh KC, Harich KC, Yu S, Lin J, Sandoe JL, Okumoto S et al: Tobacco nicotine uptake permease (NUP1) affects alkaloid metabolism. Proc Natl Acad Sci U S A. 2011, 108(44):18179–18184.

Kajikawa M, Hirai N, Hashimoto T: A PIP-family protein is required for biosynthesis of tobacco alkaloids. Plant Mol. Biol. 2009, 69(3):287–298.

Kajikawa M, Sierro N, Hashimoto T, Shoji T: A model for evolution and regulation of nicotine biosynthesis regulon in tobacco. Plant Signaling & Behavior 2017, 12(6): e1338225.

Kajikawa M, Sierro N, Kawaguchi H, Bakaher N, Ivanov NV, Hashimoto T, Shoji T: Genomic insights into the evolution of the nicotine biosynthesis pathway in tobacco. Plant Physiol. 2017, 174(2):999–1011.

Kato K, Shitan N, Shoji T, Hashimoto T: Tobacco NUP1 transports both tobacco alkaloids and vitamin B6. Phytochemistry 2015, 113:33–40.

Kazan K, Manners JM: Jasmonate signaling: toward an integrated view. Plant Physiol. 2008, 146(4):1459–1468.

Kazan K, Manners JM: MYC2: the master in action. Mol Plant 2013, 6(3):686–703.

Legg PD, Chaplin JF, Collins GB: Inheritance of percent total alkaloids in *Nicotiana tabacum* L. J. Hered. 1969, 60(4):213–217.

Legg PD, Collins GB: Inheritance of per cent total alkaloids in *Nicotiana tabacum* L. II. Genetic effects of two loci in Burley 21× LA Burley 21 populations. Can. J. Genet. Cytol. 1971, 13(2):287–291.

Michelmore RW, Paran I and Kesseli RV: Identification of markers linked to disease-resistance genes by bulked segregant analysis: a rapid method to detect markers in specific genomic regions by using segregating populations. Proc Natl Acad Sci U S A. 1991, 88(21): 9828–9832.

Miller MA, Zachary JF: Mechanisms and morphology of cellular injury, adaptation, and death, Pathologic Basis of Veterinary Disease (Sixth Edition), 2017, 2–43.

Nützmann HW, Doerr D, Ramírez-Colmenero A, Sotelo-Fonseca JE, Wegel E, Stefano MD, Wingett SW, Fraser P, Hurst L, Fernandez-Valverde SL, Osbourn A: Active and repressed biosynthetic gene clusters have spatially distinct chromosome states. Proc Natl Acad Sci U S A. 2020, doi: 10.1073/pnas.1920474117.

Nützmann HW, Huang A, Osbourn A: Plant metabolic clusters - from genetics to genomics. New Phytol. 2016, 211: 771–789.

Paul P, Singh SK, Patra P, Liu X, Pattanaik S, Yuan L: Mutually regulated AP2/ERF gene clusters modulate biosynthesis of specialized metabolites in plants. Plant Physiol. 2020, 182:840–856.

Qin Y, Bai S, Li W, Sun T, Galbraith, Yang Z, Zhou Y, Sun G, Wang B: Transcriptome analysis reveals key genes involved in the regulation of nicotine biosynthesis at early time points after topping in tobacco (*Nicotiana tabacum* L.). BMC Plant Biol. 2020, 20:30.

Reed DG, Jelesko JG: The *A* and *B* loci of *Nicotiana tabacum* have non-equivalent effects on the mRNA levels of four alkaloid biosynthesis genes. Plant Sci. 2004, 167(5): 1123–1130.

Rushton PJ, Bokowiec MT, Han S, Zhang H, Brannock JF, Chen X, Laudeman TW, Timko MP: Tobacco transcription factors: novel insights into transcriptional regulation in the Solanaceae. Plant Physiol. 2008, 147(1):280–295.

Ryan SM, Cane KA, De Boer KD, Sinclair SJ, Brimblecombe R, Hamill JD: Structure and expression of the quinolinate phosphoribosyltransferase (*QPT*) gene family in *Nicotiana*. Plant Sci. 2012, 188-189:102–110.

Sears MT, Zhang H, Rushton PJ, Wu M, Han S, Spano AJ, Timko MP: *NtERF32*: a non-*NIC2* locus *AP2/ERF* transcription factor required in jasmonate-inducible nicotine biosynthesis in tobacco. Plant Mol. Biol. 2014, 84(1-2):49–66.

Shen Q, Lu X, Yan TX, Fu XQ, Lv ZY, Zhang FY, Pan QF, Wang GF, Sun XF, Tang KX: The jasmonate-responsive AaMYC2 transcription factor positively regulates artemisinin biosynthesis in *Artemisia annua*. New Phytol. 2016, 210(4):1269–1281.

Shoji T, Harshimoto T: Tobacco MYC2 regulates jasmonate-inducible nicotine biosynthesis genes directly and by way of the *NIC2*-locus *ERF* genes. Plant Cell Physiol. 2011, 52(6):1117–1130.

Shoji T, Hashimoto T: DNA-binding and transcriptional activation properties of tobacco *NIC2*-locus *ERF189* and related transcription factors. Plant Biotechnol 2012, 29(1):35–42.

Shoji T, Hashimoto T: Nicotine biosynthesis. In plant metabolism and biotechnology. John Wiley & Sons, Ltd; 2011, 191–216.

Shoji T, Hashimoto T: Recruitment of a duplicated primary metabolism gene into the nicotine biosynthesis regulation in tobacco. Plant J. 2011, 67(6): 949–959.

Shoji T, Hashimoto T: Recruitment of a duplicated primary metabolism gene into the nicotine biosynthesis regulon in tobacco. Plant J. 2011b, 67: 949–959.

Shoji T, Hashimoto T: Smoking out the masters: transcriptional regulators for nicotine biosynthesis. Plant Biotechnol. 2013, 30: 217–224.

Shoji T, Hashimoto T: Stress-induced expression of NICOTINE 2-locus genes and their homologs encoding Ethylene Response Factor transcription factors in tobacco. Phytochem 2015, 113: 41–49.

Shoji T, Hashimoto T: Tobacco MYC2 regulates jasmonate inducible nicotine biosynthesis genes directly and by way of the *NIC2*-locus *ERF* genes. Plant Cell Physiol. 2011c, 52: 1117–1130.

Shoji T, Inai K, Yazaki Y, Sato Y, Takase H, Shitan N, Yazaki K, Goto Y, Toyooka K, Matsuoka K et al: Multidrug and toxic compound extrusion-type transporters implicated in vacuolar sequestration of nicotine in tobacco roots. Plant Physiol. 2009, 149(2):708–718.

Shoji T, Kajikawa M, Hashimoto T: Clustered transcription factor genes regulate nicotine biosynthesis in tobacco. Plant Cell 2010, 22(10):3390–3409.

Shoji T, Ogawa T, Hashimoto T: Jasmonate-induced nicotine formation in tobacco is mediated by tobacco COI1 and JAZ genes. Plant Cell Physiol. 2008, 49(7):1003–1012.

Sierro N, Battey JN, Ouadi S, Bakaher N, Bovet L, Willig A, Goepfert S, Peitsch MC, Ivanov NV: The tobacco genome sequence and its comparison with those of tomato and potato. Nat Commun 2014, 5:3833.

Sinclair SJ, Murphy KJ, Birch CD, Hamill JD: Molecular characterization of quinolinate phosphoribosyltransferase (QPRtase) in *Nicotiana*. Plant Mol. Biol. 2000, 44(5):603–617.

Stepp Burtin D, Michael AJ: Over-expression of arginine decarboxylase in transgenic plants, Biochem. J. 1997, 325:331e337

Steppuhn A, Gase K, Krock B, Halitschke R, Baldwin IT: Nicotine’s defensive function in nature. PLoS Biol 2004, 2:1074–1080.

Thurston R, Smith WT, Cooper BP: Alkaloid secretion by trichomes of *Nicotiana* species and resistance to aphids. Entomol Exp Appl 1966, 9:428–432.

Saunders JW, Bush LP: Nicotine biosynthetic enzyme activities in *Nicotiana tabacum* L. genotypes with different alkaloid levels. Plant Physiol. 1979, 64:236–240.

Todd AT, Liu E, Polvi SL, Pammett RT, Page JE: A functional genomics screen identifies diverse transcription factors that regulate alkaloid biosynthesis in *Nicotiana benthamiana*. Plant J. 2010, 62(4):589–600.

Tong Z, Xiao B, Jiao F, Fang D, Zeng J, Wu X, Chen X, Yang J, Li Y: Large-scale development of SSR markers in tobacco and construction of a linkage map in flue-cured tobacco. Breeding Science 2016, 66: 381–390.

Van Moerkercke A, Steensma P, Schweizer F, Pollier J, Gariboldi I, Payne R, Bossche RV, Miettinen K, Espoz J, Purnama PC, Kellner F, Seppanen-Laakso T, O’Connor SE, Rischer H, Memelink J, Goossens A: The bHLH transcription factor BIS1 controls the iridoid branch of the monoterpenoid indole alkaloid pathway in *Catharanthus roseus*. Proc Natl Acad Sci U S A. 2015, 112: 8130–8135.

Van Ooijen, JW: JoinMap 4.0: Software for the calculation of genetic linkage maps in experimental populations, Kyazma BV, Wageningen 2006

Wang B, Lewis RS, Shi J, Song Z, Gao Y, Li W, Chen H, Qu R: Genetic factors for enhancement of nicotine levels in cultivated tobacco. Sci Rep 2015, 5:17360.

Weigel D, Glazebrook J: Transformation of agrobacterium using the freeze-thaw method. CSH Protoc. 2006, 2006(7): pii: pdb.prot4666.

Wickham H: ggplot2: elegant graphics for data analysis. Springer 2009, New York.

Xing HL, Dong L, Wang ZP, Zhang HY, Han CY, Liu B, Wang XC, Chen QJ: A CRISPR/Cas9 toolkit for multiplex genome editing in plants. BMC Plant Biol. 2014, 14(1):327.

Xu S, Brockmoeller T, Navarro-Quezada A, Kuhl H, Gase K, Ling Z, Zhou W, Kreitzer C, Stanke M, Tang H et al: Wild tobacco genomes reveal the evolution of nicotine biosynthesis. Proc Natl Acad Sci U S A. 2017, 114(23):6133–6138.

Yang Y, Yan P, Yi C, Li W, Chai Y, Fei L, Gao P, Zhao H, Wang Y, Timko MP, et al: Transcriptome-wide analysis of jasmonate-treated BY-2 cells reveals new transcriptional regulators associated with alkaloid formation in tobacco. J. Plant Physiol. 2017, 215:1–10.

Zhang H, Hedhili S, Montiel G, Zhang Y, Chatel G, Pré M, Gantet P, Memelink J: The basic helix-loop-helix transcription factor CrMYC2 controls the jasmonate-responsive expression of the ORCA genes that regulate alkaloid biosynthesis in Catharanthus roseus. Plant J. 2011, 67:61–711.

Zhang HB, Bokowiec MT, Rushton PJ, Han SC, Timko MP: Tobacco transcription factors *NtMYC2a* and *NtMYC2b* form nuclear complexes with the *NtJAZ1* repressor and regulate multiple jasmonate-inducible steps in nicotine biosynthesis. Mol Plant 2012, 5(1):73–84.

Zhou M, Memelink J: Jasmonate-responsive transcription factors regulating plant secondary metabolism. Biotechnol. Adv. 2016, 34(4):441–449.

